# Intermolecular disulfide-bond formation in human FICD modulates the activity of the hyperactive E234G mutant

**DOI:** 10.1101/126516

**Authors:** Raffaella Magnoni, Minttu S. Virolainen, Celeste M. Hackney, Cecilie L. Søltoft, Ana P. Cordeiro, Yun Liu, Carsten Scavenius, Jan J. Enghild, Brian Christensen, James Paton, Adrienne W. Paton, Esben S. Sørensen, Lars Ellgaard

## Abstract

Endoplasmic reticulum (ER) stress that leads to the accumulation of misfolded proteins in the ER initiates the unfolded protein response (UPR). This homeostatic response activates signaling pathways that seek to reinstate a proper ER protein folding balance or induce apoptosis if ER stress persists. Recently, we and others identified human FICD (Filamentation induced by cyclic AMP domain-containing protein), an enzyme with adenylyltransferase (aka AMPylation) activity, as a new UPR target. Here, we demonstrate that FICD is functionally linked to the UPR, as evidenced by the finding that the adenylyltransferase activity of the protein induces ER stress, while FICD silencing increases sensitivity to ER stress. We identify BiP, an abundant ER chaperone and key regulator of the UPR, as the main substrate of FICD AMPylation in ER-derived microsomes, further emphasizing close functional connection of FICD to the UPR and in line with recent reports that AMPylation inactivates BiP. Notably, BiP overexpression increased the levels of BiP AMPylation as well as FICD auto-AMPylation, suggesting a homeostatic response that balances the pool of active BiP to modulate its functions in protein folding as well as UPR signaling. Finally, we show that overexpressed FICD forms a disulfide-bonded homo-dimer through Cys51 and Cys75 and demonstrate that mutation of these two cysteines in the context of a hyperactive FICD mutant leads to increased BiP AMPylation. This latter finding opens up the possibility that FICD activity is redox regulated and closely connected with ER redox homeostasis.

## Introduction

Stress conditions that perturb endoplasmic reticulum (ER) homeostasis often result in the accumulation of misfolded proteins in this organelle. This situation initiates an intricate and coordinated transcriptional and translational program known as the unfolded protein response (UPR) [1], which can also be induced by ER lipid disequilibrium [2]. ER stress is detected by three UPR sensors that all localize to the ER membrane: Protein kinase RNA-like ER kinase (PERK), Inositol-requiring protein 1 (IRE1) and Activating transcription factor 6 (ATF6). These proximal UPR transducers contain a single membrane-spanning *α*-helix that separates an ER luminal sensor domain from a cytosolic effector domain. The cellular measures initiated by the UPR collectively aim to restore ER folding capacity, e.g. by inhibiting protein translation to relieve the protein folding burden on the ER [3], degrading a subgroup of mRNAs associated with the ER membrane [4], and increasing the production of ER chaperones as well as ER-associated degradation factors that help degrade misfolded ER proteins [5-7]. Persistent ER stress resulting from an inability of the cell to restore homeostatic conditions will prompt cells to enter apoptosis.

The abundant ER protein BiP, a chaperone of the Hsp70 family, has important functions in ER protein folding and in targeting misfolded proteins for ER-associated degradation [8]. It recognizes substrates through a substrate-binding domain, while a nucleotide-binding domain controls substrate binding and release through cycles of ATP binding and hydrolysis. Co-chaperones of the Hsp40 family – seven of which (ERdj1-7) are expressed in the ER [9] – assist BiP in substrate recognition and stimulate BiP’s ATPase activity (see e.g. [10-13]). BiP also plays a crucial role in modulating the activity of PERK, IRE1 and ATF6. Under non-stressed conditions, BiP binds each of these three proteins but dissociates during ER stress to assist protein folding (reviewed in [1, 14]). This dissociation is a prerequisite for activation of the UPR transducers [15].

Previously, we have shown that filamentation induced by cyclic AMP domain-containing protein (FICD; also referred to as HYPE for “Huntingtin associated protein E”) is upregulated by conditions of oxidative stress in the ER that result in a mild UPR [16]. Additional studies have also shown FICD [17-19] and *Drosophila* FICD (*d*FIC) [20] to be targets of the UPR. FICD is the only mammalian protein with a so-called Fic domain, which uses ATP as a substrate to adenylylate (AMPylate) substrates on Ser, Thr and Tyr residues. The Fic domain is predicted to occur in almost 3000 proteins (most of them bacterial), which are involved in a variety of cellular processes (reviewed in [21, 22]). For instance, some pathogenic bacteria inject Fic domain-containing proteins into the target cell cytosol where they AMPylate Rho family GTPases, which blocks the interaction of the GTPases with downstream effectors and prevents actin polymerization, leading to collapse of the actin cytoskeleton [23].

FICD contains a single predicted transmembrane helix (residues 24-44), two tetratricopeptide repeat (TPR) motifs (residues 105-170), and a Fic domain (residues 215-432) (Fig. 1A). A TPR motif is a 34 amino acid repeat that forms a helix-loop-helix structure [24]. Most often TPR motifs are found in tandem arrays of three or more repeats that mediate protein-protein interactions [24]. The crystal structure of a fragment of human FICD (residues 103-432) containing the TPR motifs and the Fic domain [25] shows that the TPR motifs are connected to the Fic domain by a single long *α*-helix. The Fic domain was crystallized in the absence and presence of different cofactors [25], e.g. ADP and ATP, and displays known key features of this type of domain (reviewed in [21]). These include the catalytic loop bridging *α*-helices 3 and 4 that contains a characteristic catalytic motif harboring a critical His (His363 in FICD), as well as an auto-inhibitory helix (Fig. 1A). This helix contains an inhibitory motif including a crucial glutamate residue (Glu234 in FICD) that competes with the γ-phosphate of ATP for binding to the enzyme [26]. In accordance with results obtained for other Fic-domain containing proteins, the H363A and E234G mutants of FICD are catalytically inactive and hyperactive, respectively [17, 25-28]. Moreover, the E234G mutant displays a strongly increased level of auto-AMPylation [17, 25, 26, 29].

**Fig. 1.**
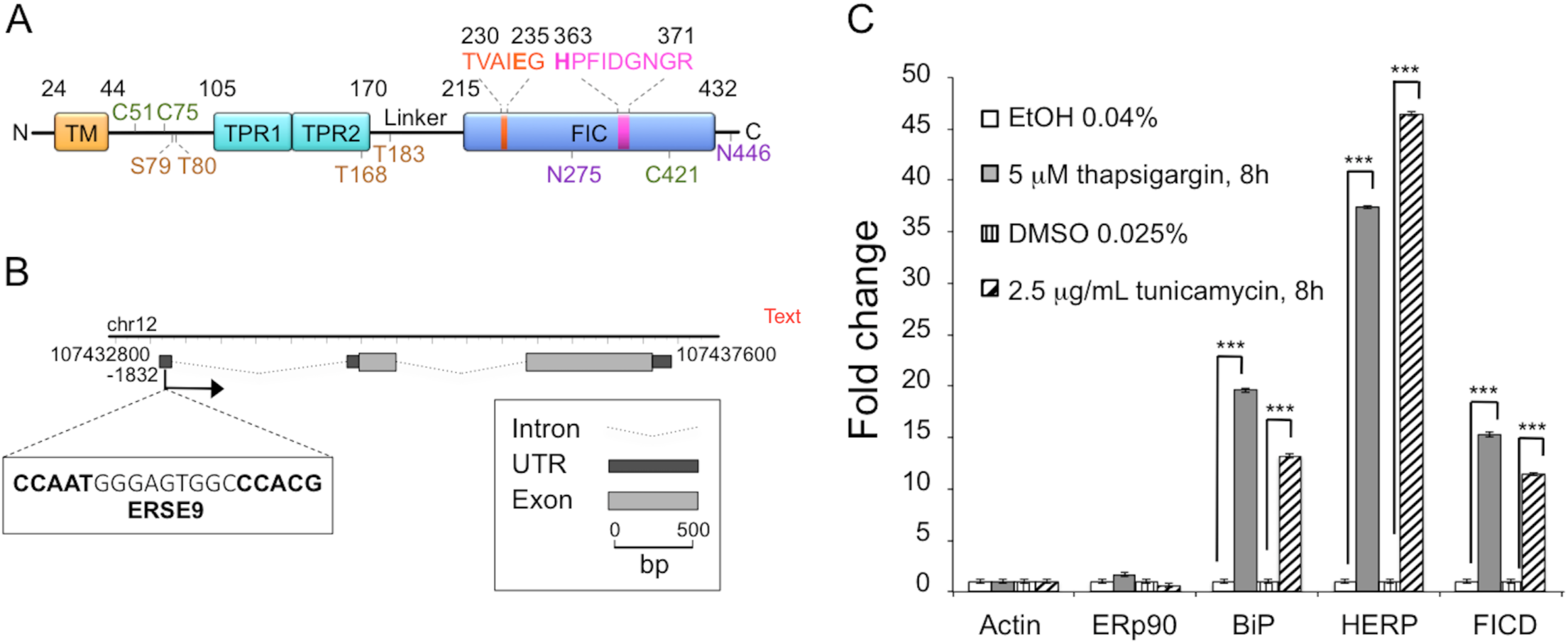
FICD expression is induced by the UPR. (A) Schematic representation of FICD. TM, transmembrane domain (residues 24-44); TPR, Tetratricopeptide repeat region (TPR1: residues 105-135, TPR2: residues 140-170); Fic, Filamentation induced by cyclic AMP domain (residues 215-432). Residues ^230^TVAIEG^235^ constitute the auto-inhibitory motif and residues ^363^HPFIDGNGR^371^ represent the active site motif. N-glycosylation sites are shown in purple (residues N275 and N446), cysteine residues are shown in green (C51, C75 and C421), and auto-AMPylation sites in orange (S79, T80, T168, T183). (B) Chromatin immunoprecipitation sequencing data obtained from the University of California–Santa Cruz Genome Browser indicate the presence of a single promoter region at position –1832. This region contains a typical ERSE9 consensus sequence (CCAAT-N9-CCACG) as shown in the figure. The FICD gene structure including untranslated regions (UTRs), introns and exons was visualized using FancyGene [41]. (C) Relative abundance of FICD mRNA in HEK293 cells analyzed by qRT-PCR. Cells were treated for 8 h with solvent (0.04% ethanol or 0.025% DMSO) or with the ER stress inducers thapsigargin (5 μΗ in 0.04% ethanol) or tunicamycin (2.5 μg/ml in 0.025% DMSO). mRNA levels were normalized to β-actin. ERp90 served as a negative control (i.e. a gene not responsive to ER stress), and BiP and HERP were positive controls. Data represent the mean of 3 biological replicates analyzed in 3 technical replicates ± SEM. Statistical analysis was performed using Student’s unpaired *t* test (two tailed, heteroschedastic).

Data have shown that several histones and cytosolic heat shock proteins are excellent *in vitro* substrates of AMPylation by hyperactive FICD and the *C. elegans* ortholog, FIC-1 [30-33], and translation elongation factors have been identified as AMPylation targets in mammalian cell extracts [28] and *C. elegans* [33]. However, BiP has emerged as an important cellular substrate of human FICD. For instance, human FICD was shown to immunoprecipitate BiP [17], and recombinant human FICD and *d*FIC are able to AMPylate both recombinant BiP [17] as well as BiP present in S2 [20] and HEK293 whole cell lysates [28]. Importantly, FICD was demonstrated to also AMPylate BiP in cells [27], a modification previously ascribed to ADP-ribosylation [34] and known to decrease at high protein load (i.e. during the UPR) to activate the protein [34-36]. Accordingly, BiP AMPylation is high at steady state (low protein load) and results in inactivation by weakening substrate interactions and inhibiting ATPase stimulation by the ERdj6 co-chaperone [27]. The same overall conclusions were reached for the *Drosophila* system [20]. An important cellular function of FICD therefore seems to be AMPylation of BiP under normal cellular conditions to generate a pool of dormant BiP for rapid reactivation during ER stress. While it is now clear that FICD also functions to de-AMPylate BiP [37], the mechanism that regulates the opposing enzymatic activities of FICD is unknown.

Here we investigate in human cells how FICD influences BiP AMPylation in response to cellular levels of BiP and upon abrogation of FICD dimerization through mutation of cysteine residues involved in intermolecular disulfide-bond formation. Our findings reveal new potential regulatory features of the interplay between FICD and BiP, and are discussed in terms of their implications for the cell biology and structure-function relations of FICD.

## Materials and Methods

### Plasmids and primers

The human FICD cDNA clone BC001342 (IMAGE clone ID 3462741) was obtained from Source BioScience Lifescience (Nottingham, U.K.). Constructs were made by the cloning of PCR products into the pcDNA3 vector (Invitrogen) using BamHI and NotI restriction sites. The correct sequences of all generated plasmids were verified by DNA sequencing. For the pcDNA3/FICD-HA_WT_ construct encoding the wild-type protein, the FICD cDNA was amplified using the HA-FICD-fwd and HA-FICD-rev primers (Table 1 provides an overview of all primers used in the present study). The resulting construct, pcDNA3/FICD-HA_WT_, was used as a template for QuikChange mutagenesis (Stratagene) to generate the pcDNA3/FICD-HA_E234G_, pcDNA3/FICD-HA_H363A_, pcDNA3/FICD-HA_C51A_, pcDNA3/FICD-HA_C75A_, and pcDNA3/FICD-HA_C421A_ mutants using the FICD-HA_E234G_-fwd and FICD-HA_E234G_-rev primers, FICD-HA_H363A_-fwd and FICD-HA_H363A_-rev primers, FICD-HA_C51A_-fwd and FICD-HA_C51A_-rev primers, FICD-HA_C75A_-fwd and FICD-HA_C75A_-rev primers and FICD-HA_C421A_-fwd and FICD-HA_C421A_-rev primers, respectively. The pcDNA3/FICD-HA_C51A_ construct was used as template for QuikChange mutagenesis to generate the pcDNA3/FICD-HA_C51A/C75A_ double mutant with the FICD-HA_C75A_-fwd and FICD-HA_C75A_-rev primers.

**Table 1.**
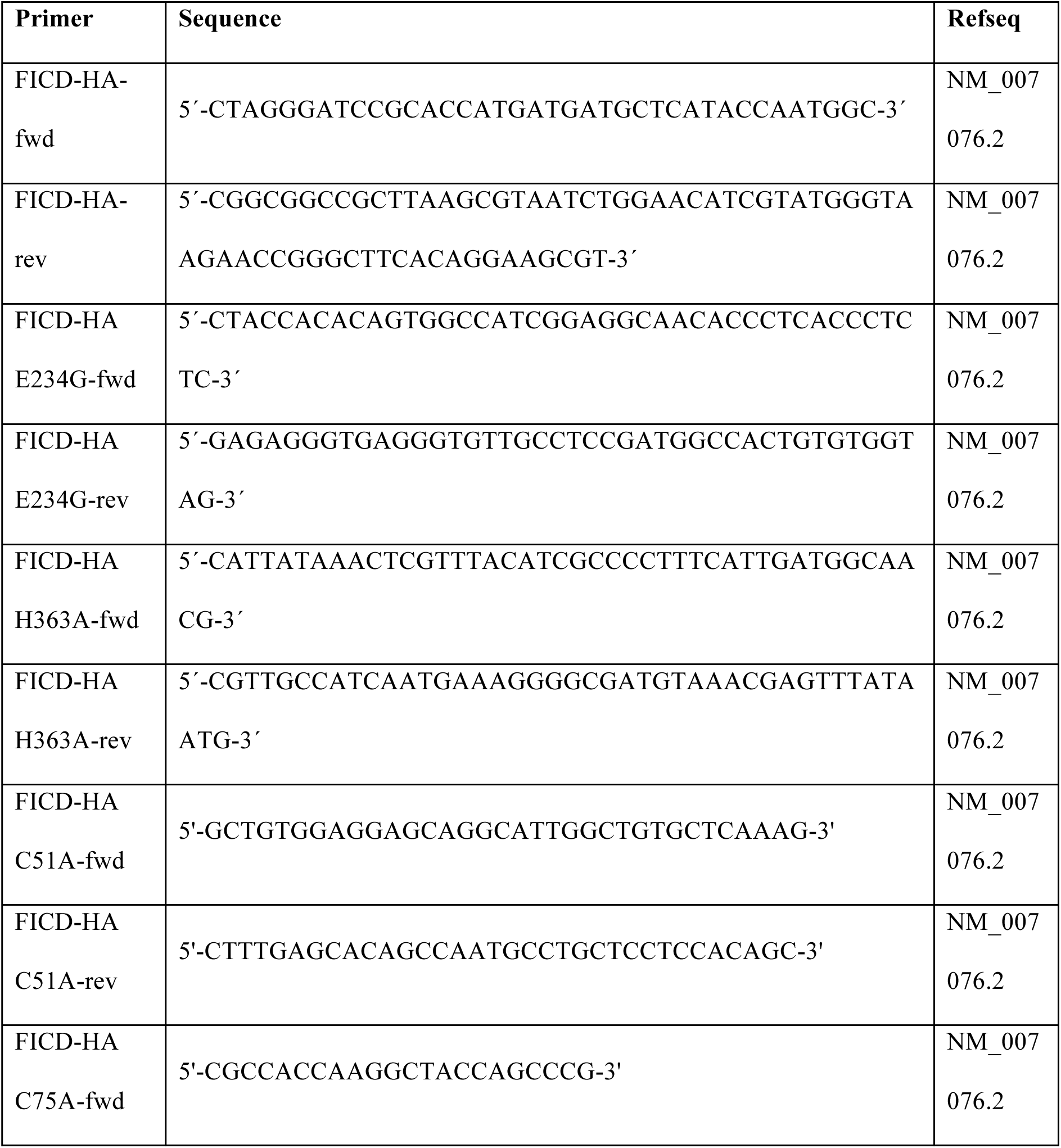
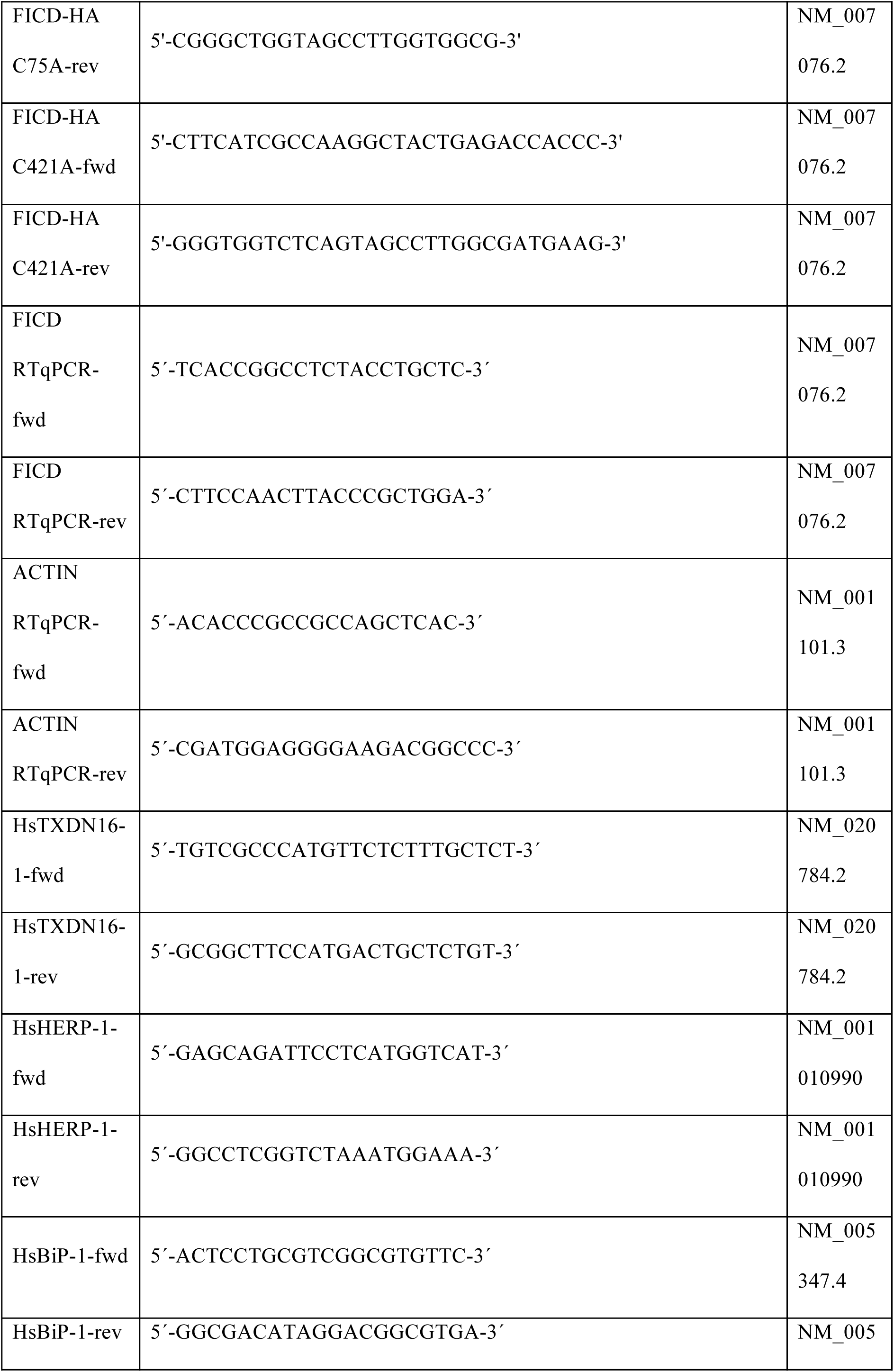
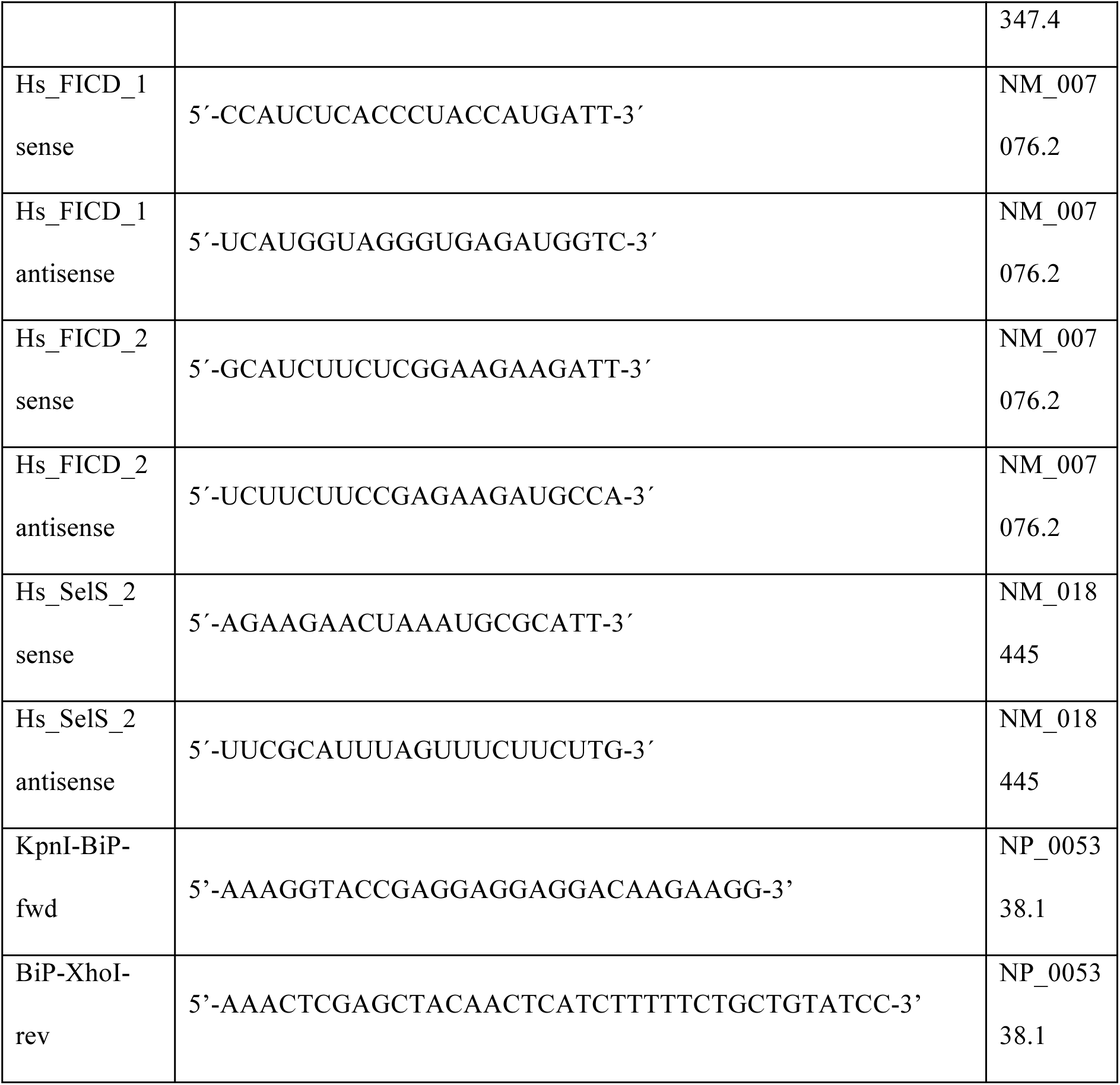
Overview of oligonucleotide primer sequences.

For generating the pcDNA3-ERp44ss-HA-BiP construct, the human BiP sequence was amplified using pCMV BiP-Myc-KDEL-WT [38] (a gift from Ron Prywes; Addgene plasmid #27164) as template and the KpnI-BiP fwd and BiP-XhoI-rev primers (see Table 1). The PCR product was then cloned into the pcDNA3–ERp44ss-HA vector [39] to generate a construct encoding the signal sequence of human ERp44 followed by HA-tagged full-length human BiP. The correct sequence of the construct was verified by DNA sequencing.

### Cell culture and transfection

Human embryonic kidney (HEK293) cells were maintained in *α*-minimal essential medium (*α*-MEM; Invitrogen), supplemented with 10% fetal bovine serum (FBS; Sigma Aldrich) at 37°C with 5% CO_2_.Transient transfection of cells was performed using polyethylenimine (PEI). Briefly, PEI (Polysciences) was dissolved in water and neutralized with 1 N HCl to a final concentration of 1mg/ml and filtered through a 0.22 μm filter. For transfection performed in one well of a 6-well plate, 2 μg plasmid DNA and 8 μl of PEI were mixed in 100 μl Opti-MEM medium (Gibco) for 10 minutes before addition to the cells. Unless otherwise stated, cells were analyzed 18 h post-transfection. For ER stress induction, untransfected HEK293 cells were treated with either 5 μM thapsigargin (Sigma) in 0.04% ethanol or 2.5 μg/ml tunicamycin (Sigma) in 0.025% dimethyl sulfoxide (DMSO) for 8 h. Control cells for thapsigargin and tunicamycin treatments were incubated with 0.04% ethanol or 0.025% DMSO, respectively.

### Quantitative real-time (qRT)-PCR

RNA was extracted from cells using TRI Reagent (Sigma). cDNA was synthetized by the RevertAid Premium Reverse Transcriptase (Fermentas) using poly-dT primers. Real-time PCR reactions were performed on a CFX96 Real-time PCR Detection system (Bio-Rad) using the Power SYBR Green PCR Master Mix (Applied Biosystems). qRT-PCR reaction efficiencies were analyzed and threshold cycle values were used to assess the relative expression levels normalized to actin or HRPD1 using the 2(-ΔΔCt) method. ERp90 was used as negative control as the expression of the gene is unaffected by ER stress [16], whereas BiP and HERP were used as positive controls for UPR target genes. Gene-specific primers are provided in Table 1. Data represent the mean of 3 biological replicates analyzed in 3 technical replicates ± SEM. Statistical analysis was performed using Student’s unpaired *t* test (two tailed, heteroschedastic).

### Computational analyses

Chromatin immunoprecipitation sequencing (ChIP-seq), CpG-island, and histone modification data were obtained from the University of California–Santa Cruz Genome Browser (http://genome.ucsc.edu; [40]). Further sequence analysis was carried out using CLC Main Workbench software (CLCBio, Aarhus, Denmark). The FICD gene structure including UTRs, introns and exons was visualized using FancyGene (http://bio.ieo.eu/fancygene/; [41]).

### Antibodies

The following mouse monoclonal antibodies were used: anti-LDH (Lactate dehydrogenase) (SC-133123, Santa Cruz Biotechnology), anti-GFP (A6455, Invitrogen), anti-HA (12CA5, Roche; HA.11, Covance), and anti-β-actin (AC-15, Sigma). The rabbit polyclonal antisera used were: anti-BiP (G8918, Sigma), anti-ERp57 (a gift from A. Helenius, Zürich, Switzerland), anti-HERP (a gift from L. Hendershot, Memphis, TN), anti-TMX3 [42], anti-Giantin (ab24586, Abcam), and anti-CD4 (H-370, Santa Cruz Biotechnology).

### Gel electrophoresis and Western blotting

Transfected HEK293 cells were washed and incubated with 20 mM N-ethylmaleimide (NEM) (Sigma) in PBS (phosphate-buffered saline; 154 mM NaCl, 1.9 mM KH_2_PO_4_, 8.1 mM Na_2_HPO_4_, pH 7.3, 1 mM CaCl_2_, 0.5 mM MgCl_2_), 20 min on ice. Cells were scraped off in lysis buffer (80 mM Tris-HCl, pH 6.8, 1% SDS, 200 μM phenylmethanesulfonylfluoride (PMSF)). Samples for Western blotting were boiled in SDS-PAGE loading buffer (25 mM Tris-HCl, pH 6.8, 1% SDS, 10% glycerol, 0.002% bromophenol blue) without reducing agents, unless otherwise stated. Samples were separated by SDS-PAGE on Tris-glycine polyacrylamide Hoeffer minigels (GE Healthcare) and transferred onto a polyvinylidene difluoride (PVDF) membrane. Membranes were probed with primary antibodies in the following dilutions: anti-HA, 1:1,000; anti-β-actin, 1:25,000; anti-ERp57, 1:1,000; anti-HERP, 1:2,000; anti-BiP, 1:5,000; anti-TMX3, 1:500; anti-LDH, 1:1000. After the addition of horseradish peroxidase-conjugated goat anti-rabbit or anti-mouse IgG (Pierce) secondary antibody (1:100,000), the bound antibody was detected with the ECL Select detection reagent (GE Healthcare).

### Confocal immunofluorescence microscopy

Transfected HEK293 cells on microscope coverslips were fixed 18 h post-transfection using 4% paraformaldehyde (PFA). To study ER colocalization cells were co-transfected with pcDNA3/FICD-HA_WT_ and pCMV/myc/ER/GFP (Invitrogen). To stain intracellular proteins, cells were blocked with a solution of 3% bovine serum albumin (BSA) in PBS for 15 min and treated with Permeabilization buffer (P-buffer; 0.05% saponin, 3% BSA in PBS) at room temperature, for 30 min. Next, cells were incubated with primary antibody for 1 h at room temperature in the following dilutions: anti-HA, 1:1000; anti-GFP, 1:1000; anti-Giantin, 1:1000; anti-CD4, 1:100. Cells were then washed three times in PBS and incubated for 45 min with secondary antibodies diluted 1:1000 in P-buffer. Alexa 594 goat anti-mouse and Alexa 488 goat anti-rabbit were used as secondary antibodies (Molecular Probes). Coverslips were rinsed with PBS and finally water, and mounted on slides with Gel Mount^™^ Aqueous Mounting Medium (Sigma). For immunofluorescence microscopy of cell surface exposed proteins on non-permeabilized cells, transfected cells were fixed in 4% PFA, blocked with a solution of 3% BSA in PBS, and incubated with primary and secondary antibodies diluted in PBS, avoiding the use of saponin. Nuclei were stained with the Hoechst 33342 dye (1 μM in PBS) for 5 min. Confocal images of fixed cells were acquired using a Point-scanning Confocal Microscope SP5-X MP, Leica Microsystems.

### Cell fractionation

HEK293 cells were first transfected with FICD or BiP-encoding constructs, as specified below. After 18 h cells were washed with PBS and incubated with 20 mM NEM (Sigma) in PBS for 20 min on ice. Cells were scraped off in PBS and harvested by centrifugation at 4,000 g for 5 min at 4°C. The cells were resuspended in homogenization buffer (20 mM HEPES-NaOH pH 7.5, 0.25 M sucrose, 200 μM PMSF, 1x cOmplete EDTA-free protease inhibitor cocktail (Sigma-Aldrich) and 20 mM NEM) as indicated. The cells were homogenized by passing 10 times through a 27-gauge needle. After removal of cell debris by centrifugation at 500 g for 5 min at 4°C, crude membranes were isolated through centrifugation at 100,000 g for 1h at 4°C.

### Proteinase K protection assay

To determine the topology of FICD, HEK293 cells were transfected with constructs to express FICD-HA_WT_. Crude membranes isolated as described above were resuspended in assay buffer (25 mM Tris-HCl, pH 8, 500 mM NaCl) and treated with 0.2 mg/ml Proteinase K (Roche Molecular Biochemicals) at room temperature for 20 min in the presence or absence of 1% Triton X-100. After inhibition of Proteinase K with 5 mM PMSF, samples were separated by SDS-PAGE under non-reducing conditions and proteins were visualized by Western blotting.

### Pulse-chase analysis and immunoprecipitation (IP)

8 h post-transfection, cells were washed with starvation medium (Dulbecco’s modified Eagle’s medium without methionine and cysteine from Sigma), incubated in starvation medium for 15 min at 37°C, and pulsed overnight at 37°C with 100 μCi/ml [^35^S]-methionine/cysteine (PerkinElmer). Cells were washed once in chase medium (*α*-MEM/10% FCS containing 10 mM methionine) and incubated in the same medium for 30 min at 37°C. After the chase, dishes were transferred to ice and washed with PBS. Cells were scraped off in 200 μl lysis buffer (30 mM Tris-HCl, pH 8.0, 100 mM NaCl, 5 mM EDTA, 1.5% SDS). Lysates were homogenized by sonication using a UP50H ultrasonic processor (Hielscher) (20 cycles of 0.9 s, 90% amplitude) and diluted 1:4 in denaturing IP buffer (30 mM Tris-HCl, pH 8.0, 100 mM NaCl, 5mM EDTA, 2.5% Triton X-100). After 1 h incubation on ice, samples were centrifuged at 25,000 g for 30 min at 4°C, and the supernatant was rotated overnight at 4°C with 30 μl Protein A-Sepharose beads (GE Healthcare) preadsorbed with 1 μl anti-HA antibody. After purifying the protein of interest overnight with the Protein A-bound antibodies, the supernatant was removed. The beads were washed 4 times with denaturing IP buffer + 1% Triton X-100 and one time with denaturing IP buffer without Triton X-100. The supernatant was removed, and 12 μl SDS-PAGE loading buffer was added to each sample. Proteins were eluted from the beads by boiling in SDS-PAGE loading buffer.

### Endoglycosidase H digestion

Endoglycosidase H (Endo H) digestions were performed on pulse-labeled immunoprecipitates after release from the beads by denaturation at 100°C for 5 min in lysis buffer (80 mM Tris-HCl, pH 6.8, 1% SDS, 200 μM PMSF). Samples were then supplemented with 1/10 volume of NEB G5 buffer, and treated with Endo H (New England Biolabs) for 1 h at 37°C.

### RNA interference

For gene silencing mediated by RNA interference, HEK293 cells were transfected using Lipofectamine RNAiMax reagent (Invitrogen). To silence FICD, cells were transfected with two different Silencer Select (Ambion) short interfering RNAs (siRNAs) targeting FICD (Hs_FICD_1 and Hs_FICD_2) at a concentration of 5 nM. Using this procedure, the level of FICD was reduced by 80% after 24 h transfection as evaluated by qRT-PCR (Supplementary S1 Fig.). To silence Selenoprotein S/VIMP, cells were transfected with 10 nM Hs_SelS_2 siRNA from Qiagen. Gene-specific oligonucleotide sequences are provided in Table 1. As negative control, cells were transfected with 10 nM non-targeting siRNA (ntRNA) (Qiagen, 1022076).

### Metabolic activity assay

To evaluate cell viability, HEK293 cells were seeded in 96-well culture plates at a density of 1x10^3^ cells per well. After 24 h, cells were transfected with siRNA targeting FICD (Hs_FICD_1 and Hs_FICD_2,) or siRNA targeting Selenoprotein S/VIMP (Hs_SelS_2) as positive control. Cells transfected with non-targeting siRNA (Invitrogen, AM4635) were used as negative control. After 48 h of transfection cells were treated, where indicated, with 2 μg/ml tunicamycin or with 5 μM thapsigargin for 24 h to induce ER stress. As positive control for cytotoxicity, 500 mM DMSO was added for 24 h. The metabolic activity was then measured by incubating the cells with 0.4 mg/ml 3-(4,5-dimethylthiazol-2-yl)-2,5-diphenyltetrazolium bromide (MTT) (Sigma) in *α*-MEM at 37°C for 4 h. Upon removal of the medium, formazan crystals were solubilized in 200 μl of 0.04 M HCl in absolute isopropanol. The absorbance of reduced dye was measured at 540 nm. Data represent three independent experiments with 6 technical replicates in each experiment. Statistical significance (p<0.05) was assessed by performing Student’s unpaired t test (two tail, heteroschedastic).

### AMPylation assay

To determine the substrates of AMPylation by FICD in a crude membrane preparation, HEK293 cells were transfected with pcDNA3/FICD-HA_E234G_. Crude membranes isolated from transfected cells (as described above) were re-suspended in assay buffer (50 mM HEPES, 150 mM NaCl, 1 mM MgCl_2_, 200 μM PMSF, 1x cOmplete EDTA-free protease inhibitor cocktail (Sigma-Aldrich), 100 μM unlabeled (“cold”) ATP and 50 μCi [*α*-^32^P] ATP). The assay was carried out at room temperature for 1 h. After incubation, an excess of cold ATP (1 mM) was added to the membrane preparations. Next, the membranes were solubilized by adding 1% Triton X-100 (v/v) and heated to 95°C for 5 min in 1x SDS-PAGE loading buffer. Where indicated, samples were treated with 2 units of snake venom type I phosphodiesterase (PDE) for 1 h at room temperature, or 100 ng/μl subtilase cytotoxin SubAB or inactive SubA_A272_B for 1h at 30°C. When treated with SubAB, no protease inhibitors were added to the assay buffer (see above).

To study the effect of mutating Cys51 and Cys75 in FICD on AMPylation activity, the membranes were isolated from untransfected cells or HEK293 cells expressing FICD-HA_WT_, FICD-HA_C51A/C75A_, FICD-HA_E234G_ or FICD-HA_E234G/C51A/C75A_, and subjected to the AMPylation assay as described above. Following the addition of cold ATP, the crude membrane fractions were diluted in 5 μl lysis buffer (10 mM HEPES, pH 7.4, 10 mM KCl, 0.1 mM EDTA, 0.1 mM EGTA, 1% Triton X-100 (v/v), 200 μM PMSF and 1x cOmplete EDTA-free protease inhibitor cocktail (Sigma-Aldrich) and lysed for 40 min on ice, with vortexing every 10 min. The insoluble material was collected by centrifugation at 100,000 g for 60 min at 4°C. The supernatant was combined with SDS-PAGE loading buffer (reducing or non-reducing, as indicated in the figure legends) and heated to 95°C. Upon SDS-PAGE, the samples were analysed by phosphorimaging and Western blotting.

### Immunoprecipitation (IP) from crude membrane preparations

For immunoprecipitation (IP) FICD-HA_WT_, FICD-HA_H363A_, or FICD-HA_E234G_ were co-expressed with BiP-HA in HEK293 cells. The membranes were isolated and subjected to the AMPylation assay as described above. After addition of 1 mM cold ATP, membrane preparations were diluted in 500 μl of lysis buffer (10 mM HEPES, pH 7.4, 10 mM KCl, 0.1 mM EDTA, 0.1 mM EGTA, 1% Triton X-100 (v/v), 200 μM PMSF and 1x cOmplete EDTA-free protease inhibitor cocktail (Sigma-Aldrich)) and lysed by rotating at 4°C for 40 min. The insoluble material was collected by centrifugation at 100,000 g for 60 min at 4°C. 5% of the supernatant was saved as an input sample for SDS-PAGE, and the rest of the cleared lysate was combined with 40 μl of Protein A-Sepharose beads (GE Healthcare), preadsorbed with 2 μl of anti-HA antibody in lysis buffer, and rotated at 4°C overnight.

Captured proteins were harvested by centrifugation at 500 g at 4°C for 2 min. The beads were washed three times in 1 ml of lysis buffer, and proteins were eluted by adding 25 μl of loading buffer and heating to 95°C for 5 min. Proteins were separated by SDS-PAGE under non-reducing conditions and transferred onto a PVDF membrane. The PVDF membranes were first analyzed by phosphorimaging and then by Western blotting.

### Sample preparation for mass spectrometry (MS)

18 h post-transfection, HEK293 cells were washed with PBS containing 20 mM NEM, incubated in the same buffer for 20 min on ice, and scraped off in 4 ml lysis buffer (20 mM HEPES, 150 mM NaCl, 1% Triton X-100, 10% glycerol, 1 mM EDTA, 200 μM PMSF, 20 mM NEM). Cells were homogenized by passing the suspension 10 times through a 27-gauge needle. After removal of cell debris by centrifugation at 25,000 g for 30 min at 4°C, the supernatant was dialyzed overnight against 1 l of dialysis buffer (20 mM HEPES, 150 mM NaCl, 1% Triton X-100, 10% glycerol, 1 mM EDTA). The dialyzed sample was rotated overnight at 4°C with 120 μl Protein A-Sepharose beads (GE Healthcare) preadsorbed with 4 μl anti-HA antibody. The supernatant was removed and the beads were washed 5 times with wash buffer (20 mM HEPES, 150 mM NaCl, 10% glycerol, 1 mM EDTA) without Triton X-100. Proteins were eluted from the beads by incubating in 40 μl of elution buffer (6 M Urea, 1 mM EDTA, 10 mM Tris-HCl pH 6.8, 20 mM NEM) for 30 min at room temperature. Proteins were boiled in loading buffer and separated by SDS-PAGE under non-reducing conditions. The gel was either stained with Coomassie blue or colloidal Coomassie blue and the band corresponding to the FICD dimer was excised from the gel and analyzed by MS.

### MS analysis

Prior to MS analysis, the excised gel bands were in-gel digested with trypsin [43] and the tryptic peptides were micro-purified using C18 PTFE membrane disks [44]. Liquid chromatography-MS/MS analyses were performed on an EASY-nLC II system (ThermoScientific) connected to a TripleTOF 5600 mass spectrometer (AB Sciex) equipped with a NanoSpray III source (AB Sciex) operated under Analyst TF 1.5.1 control. The micro-purified sample was suspended in 0.1% formic acid, injected, trapped and desalted on a 2 cm x 100 μm trap column packed in-house with RP ReproSil-Pur C18-AQ 3 μm resin (Dr. Marisch GmbH, Ammerbuch-Entringen, Germany). The peptides were eluted from the trap column and separated on a 15-cm analytical column (75 μm i.d.) packed in-house in a pulled emitter with RP ReproSil-Pur C18-AQ 3 μm resin (Dr. Marisch GmbH, Ammerbuch-Entringen, Germany). Peptides were separated using a 20 min gradient from 5% to 35% phase B (0.1% formic acid and 90% acetonitrile) and a flow rate of 250 nl/min. For protein identification, the collected MS files were converted to Mascot generic format (MGF) using the AB SCIEX MS Data Converter beta 1.1 (AB SCIEX) and the “proteinpilot MGF” parameters. The generated peak lists were searched against the Swiss-Prot database (human) using the Mascot search engine (Matrix Science). Search parameters had NEM and propionamide as variable modifications of Cys, and peptide tolerance and MS/MS tolerance were set to 10 ppm and 0.1 Da respectively. Disulfide-bonded peptides were identified using Peakview 1.2 (AB Sciex) by manually inspecting and annotating MS/MS spectra with precursor masses matching the theoretical values.

## Results

### ER stress induces FICD expression

Constitutive overexpression of a hyperactive mutant of Ero1*α*, an ER flavoprotein oxidase, induces a mild UPR [16]. Under these conditions, we previously showed that FICD expression is significantly upregulated [16]. Employing FICD sequence analysis with the ENCODE consortium [45], we observed that the genomic data predict FICD to have one single promoter region characterized by the presence of a putative CpG island, but no TATA-box. Notably, we identified a region in the 5′UTR, in position −1832, that shows a typical ER stress response element (CCAAT-N(9)-CCACG) [46] (Fig. 1B). Additional potential UPR elements have recently been identified [17]. To test if the FICD transcript was upregulated by different stress conditions that induce the UPR, we treated HEK293 cells with either tunicamycin or thapsigargin, drugs known to initiate a strong UPR. We confirmed induction of ER stress by the transcriptional upregulation of two well-established UPR targets, BiP and HERP [47]. Under these conditions, we observed a significant increase in FICD transcript levels when compared to untreated HEK293 cells (Fig. 1C). These results demonstrate that FICD is a UPR target and corroborate recent findings from other studies [17, 20].

### FICD is a type II transmembrane glycoprotein of the ER

To analyze the subcellular localization of FICD we created a C-terminally HA-tagged fusion protein (FICD-HA_WT_). HEK293 cells transfected with the construct encoding FICD-HA_WT_ were analyzed by confocal immunofluorescence microscopy, using GFP-KDEL as a marker for the ER and staining for giantin and CD4 to label the Golgi and plasma membrane, respectively. As judged by the strongly overlapping signals for FICD-HA_WT_ and GFP-KDEL, FICD-HA_WT_ localized to the ER (Fig. 2).

**Fig. 2.**
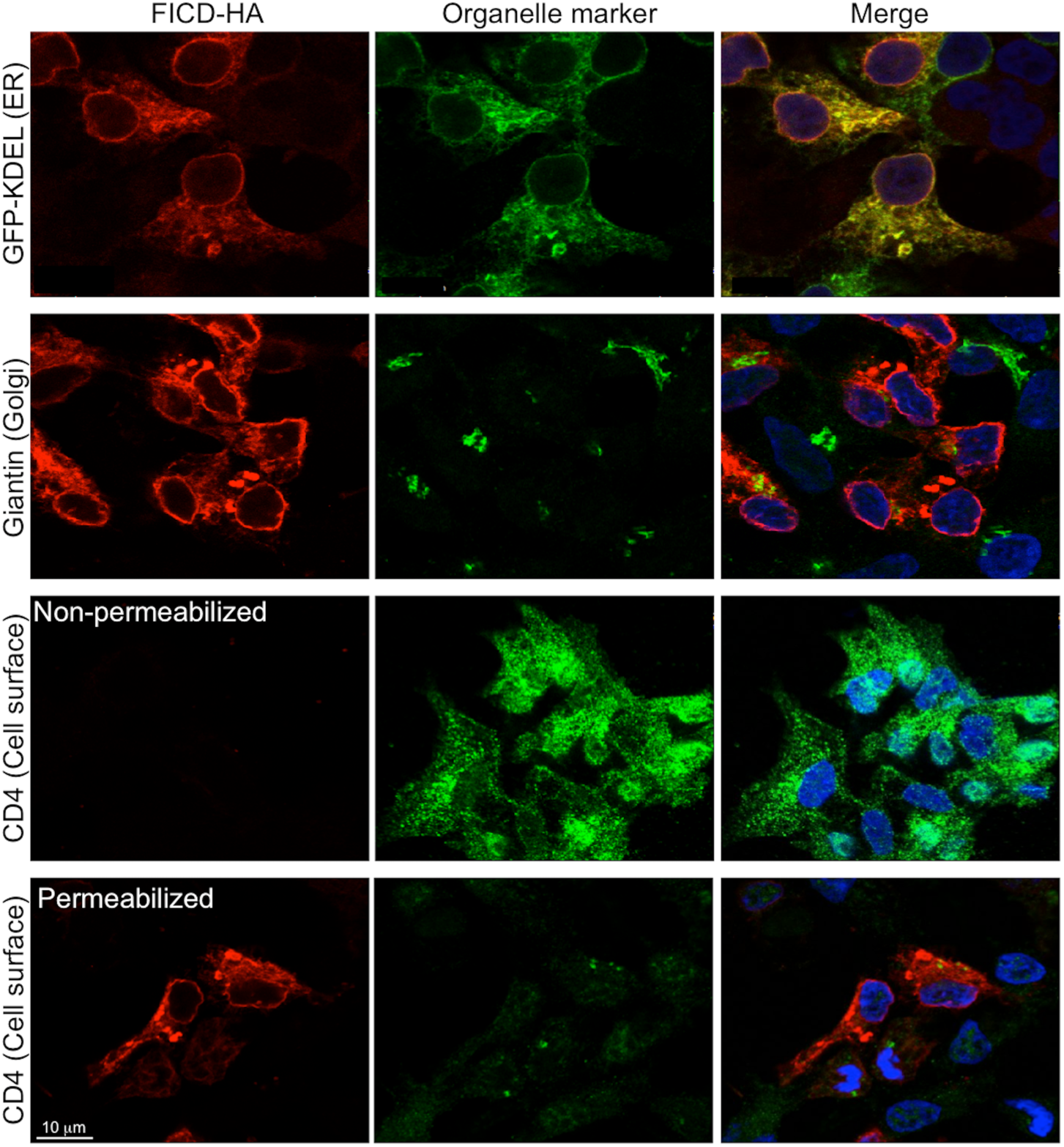
FICD localizes to the ER by confocal fluorescence microscopy. HEK293 cells expressing FICD-HA_WT_ for 18 h, stained for FICD-HA_WT_ (red) and organelle markers (green): GFP-KDEL (ER), giantin (Golgi), and CD4 (cell surface). FICD-HA_WT_ exclusively shows colocalization with the ER marker GFP-KDEL. In addition to the reticular staining pattern, a clear signal is observed at the nuclear rim. Nuclei (blue) were stained with Hoechst. Scale bar: 10 μm.

The presence of two predicted N-glycosylation sites, the conserved Asn275 as well as Asn446 (Fig. 1A), prompted us to investigate the glycosylation status of FICD-HA_WT_ expressed in HEK293 cells. The protein migrated as three distinct bands by SDS-PAGE (Fig. 3A). When treated with Endo H, an endoglycosidase that cleaves only high mannose type N-glycans, the three bands collapsed into the lower molecular weight band (Fig. 3A), indicating that FICD-HA_WT_ exists as three forms containing 0, 1 or 2 Endo H-sensitive N-glycans. Indeed, mutation of Asn275 and Asn446 abolishes FICD N-glycosylation as shown by the Mattoo lab [17].

**Fig. 3.**
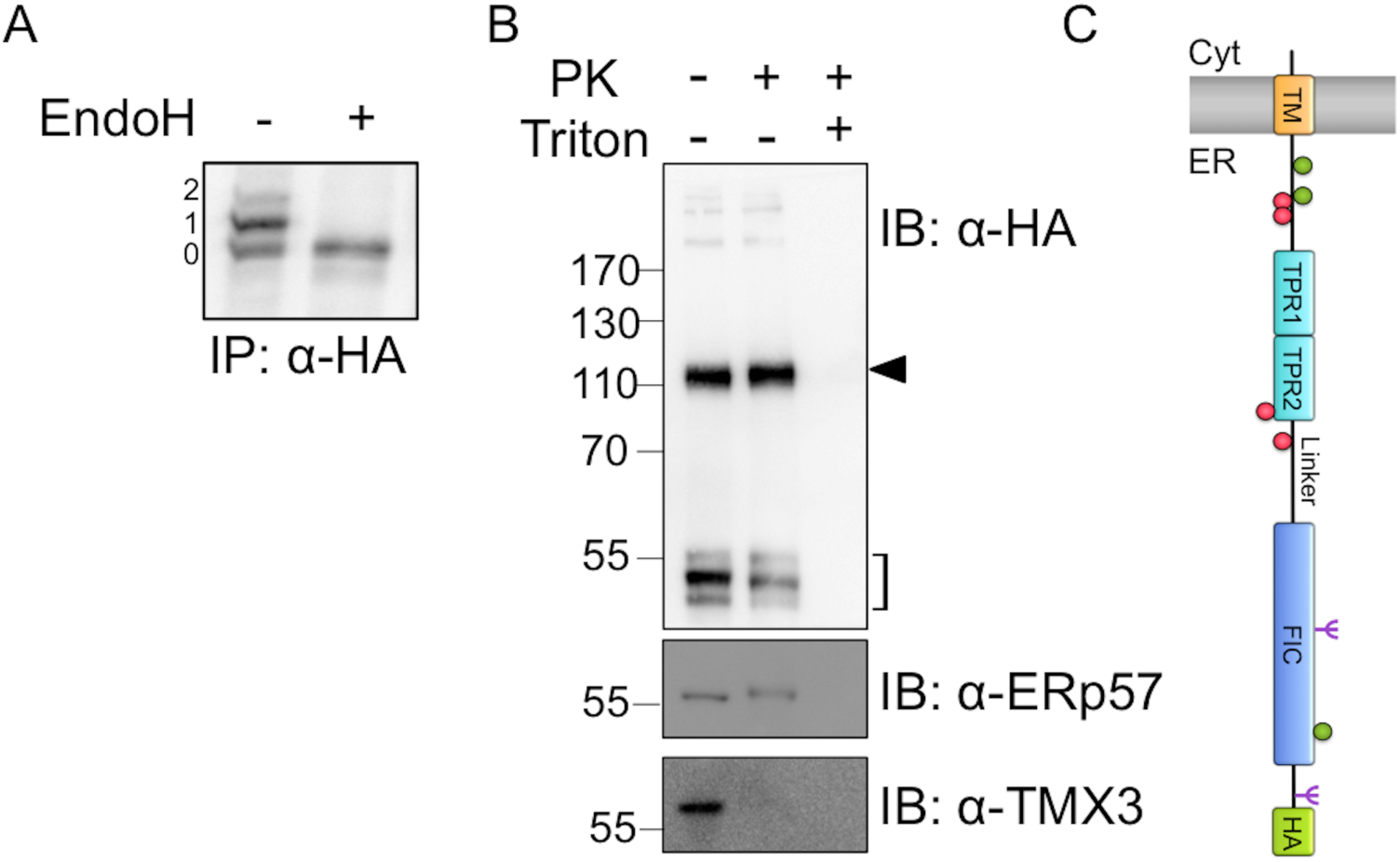
FICD is a type II transmembrane glycoprotein. (A) HEK293 cells expressing FICD-HA_WT_ were labeled with [^35^S]-methionine. Samples were immunoprecipitated with anti-HA antibody and treated with Endo H as indicated. 0, 1 and 2 refer to the number of N-glycans present on FICD-HA_WT_. (B) Determination of FICD membrane topology. Crude membranes isolated from HEK293 cells expressing FICD-HA_WT_ were treated with Proteinase K (PK) in the presence or absence of 1% Triton X-100. The square bracket indicates the FICD-HA_WT_ monomer at the predicted size of 53 kDa, and the black arrowhead the presence of a higher molecular weight complex involving FICD-HA_WT_. ERp57, a soluble protein of the ER lumen, was used to demonstrate intactness of microsomes. tmX3, a type I membrane protein of the ER, was included to show the efficiency of PK towards a protein for which the epitope faces the cytosolic side of the ER membrane. The position of molecular weight marker bands is indicated to the left of the gel in kDa. This gel was run under non-reducing conditions. (C) Schematic representation of FICD-HA_WT_ membrane topology. N-glycans are shown in purple (Ψ), cysteines as green circles, and auto-AMPylation sites as red circles. Domain coloring is given as in Fig. 1A; C-terminal HA tag is shown in green.

The presence of Endo H-sensitive glycans in the C-terminal portion of the protein strongly indicated FICD to have a type II orientation. Yet, FICD lacks an apparent arginine-based ER-retention signal typical of many type II transmembrane proteins. Therefore, we used a Proteinase K protection assay to determine the membrane topology of FICD-HA_WT_ (Fig. 3B). A crude membrane preparation containing ER-derived microsomes isolated from HEK293 cells transfected with FICD-HA_WT_ was left untreated or treated with Proteinase K in the presence or absence of Triton X-100. The untreated sample displayed the three bands at the expected size for FICD-HA_WT_ of ∼53 kDa (Fig. 3B, square bracket). Upon Proteinase K treatment in the absence of detergent, all bands corresponding to FICD-HA_WT_ were still detectable, but showed a small downward shift in migration, likely corresponding to the cleavage of at least a portion of the N-terminal 23 residues not protected by the ER membrane. Upon membrane solubilization, the signal for FICD-HA_WT_ disappeared, demonstrating that the HA-tagged C-terminus of FICD protein was protected by the ER membrane. The soluble luminal ER protein, ERp57, was used as a control for membrane integrity, and an antibody raised against the cytosolic C-terminus of TMX3, a type I ER membrane protein, was used to demonstrate efficient Proteinase K digestion of a protein for which the epitope faces the cytosolic side of the ER membrane [42]. Taken together, the data show that FICD is a type II transmembrane protein of the ER with the Fic domain facing the ER lumen as schematically depicted in Fig. 3C. Using the ER fraction of rat liver tissue and trypsin as the protease [17] or *in vitro* translation in the presence of microsomes [48], similar results have been obtained for endogenous human FICD and *d*FIC, respectively.

### FICD-HA_WT_ forms a disulfide-bonded homo-dimer involving Cys51 and Cys75

As revealed in Fig. 3B, an anti-HA reactive species appeared at∼115 kDa (black arrowhead), roughly corresponding to the size of a FICD-HA_WT_ homo-dimer. We therefore tested the sensitivity of this FICD-HA_WT_ complex to reducing conditions. While the FICD-HA_WT_ complex was clearly visible under non-reducing conditions (Fig. 4A, lane 1), it disappeared upon DTT treatment (Fig. 4A, lane 2). Moreover, no additional band appeared upon reduction suggesting that the FICD-HA_WT_ complex comprised solely two molecules of FICD-HA_WT_ held together by one or more disulfide-bonds.

**Fig. 4.**
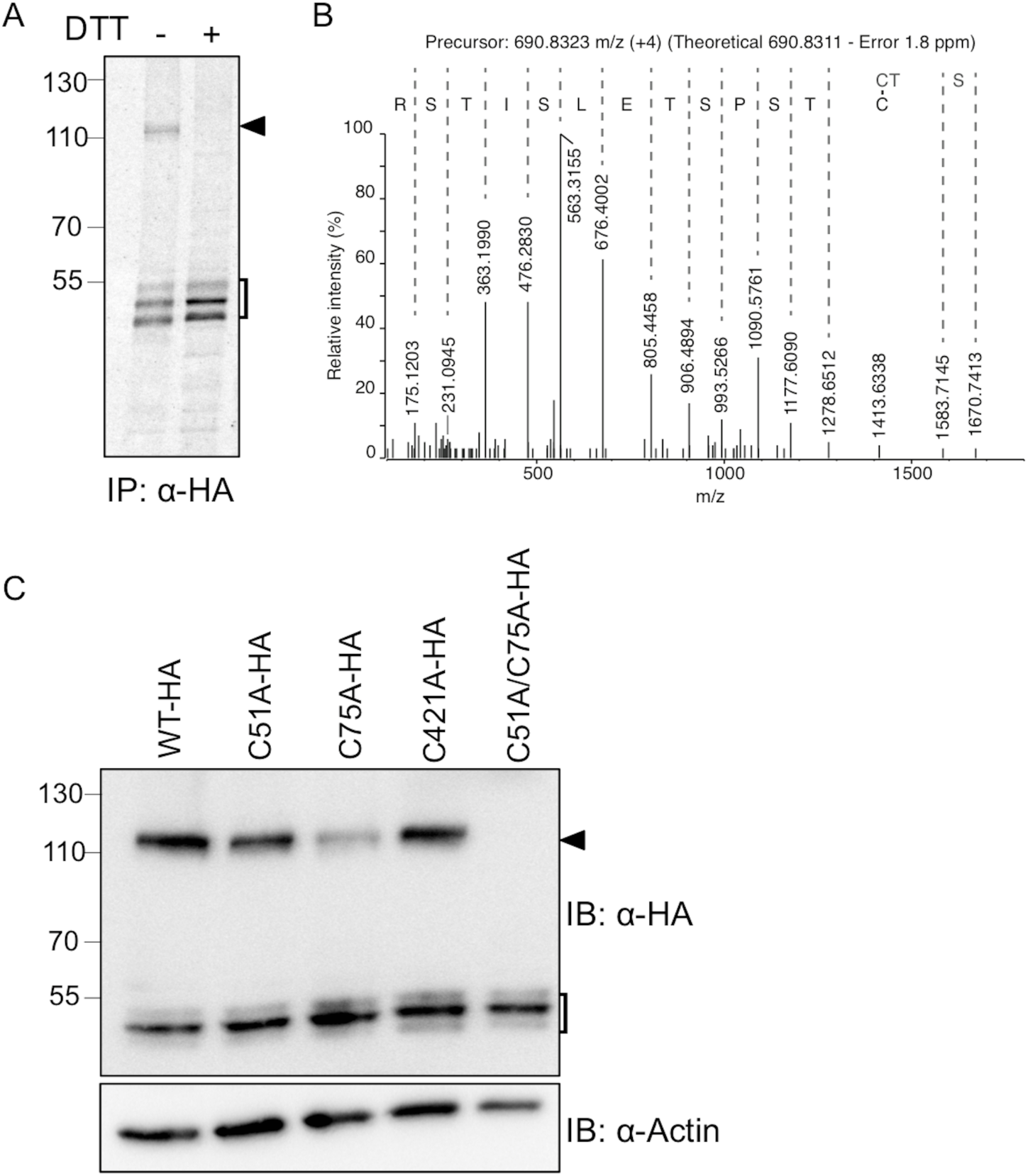
FICD-HA forms a disulfide-bonded homo-dimer involving Cys51 and Cys75. **(A)** HEK293 cells expressing FICD-HA_WT_ were labeled with [^35^S]-methionine. Samples were immunoprecipitated with anti-HA antibody and separated by SDS-PAGE under non-reducing (-) or reducing (+; 10 mM DTT) conditions. The square bracket indicates FICD-HA_WT_ monomeric forms, and the black arrowhead the disulfide-bonded dimer of FICD-HA_WT_. (B) The FICD-HA_WT_ dimer was isolated by SDS-PAGE, in-gel digested with trypsin and analyzed by LC-MS/MS. An annotated MS/MS spectrum matching the disulfide-bonded Cys75 homo-dimer is shown. The y-ion series is labeled and includes fragments covering the disulfide bond. (C) Cell were transfected with FICD-HA_WT_, FICD-HA_C51A_, FICD-HA_C75A_, FICD-HA_C421A_, or FICD-HA_C51A/C75A_ as indicated. FICD dimer formation was analyzed by Western blotting under non-reducing conditions. β-actin was used as loading control. The square bracket indicates FICD-HA monomeric forms, and the black arrowhead the disulfide-bonded dimer of FICD-HA.

To investigate this possibility, the NEM-alkylated and immunoisolated FICD-HA_WT_ complex was excised from a Coomassie-stained SDS-PAGE gel, and analyzed by LC-MS/MS. The analysis clearly identified FICD as the major constituent in the sample with a Mascot score of 4831 and an exponentially modified Protein Abundance Index (emPAI) of 13.9, more than 4 times higher than the second most abundant protein (Human C1CT). Since the Mascot search engine only identifies linear peptides, a manual search for disulfide-bonded peptides was performed. Based on the precursor mass and MS/MS spectra, we could identify the presence of a Cys75 disulfide-bonded homo-peptide (Fig. 4B). We were not able to identify any disulfide-bonded peptides involving Cys51 or Cys421, the other two cysteines in FICD.

To further investigate the involvement of Cys75, Cys51 and Cys421 in disulfide-bond formation, we generated three single point mutants, FICD-HA_C51A_, FICD-HA_C75A_ and FICD-HA_C421A_, as well as the FICD-HA_C51A/C75A_ double mutant. We observed a decrease in the amount of the dimer band for FICD-HA_C51A_ and FICD-HA_C75A_ relative to FICD-HA_WT_, whereas no difference was observed for FICD-HA_C421A_ (Fig. 4C). Western blot analysis of the double mutant FICD-HA_C51A/C75A_ showed complete loss of the dimer (Fig. 4C, lane 5), demonstrating the involvement of both Cys51 and Cys75 in the formation of the disulfide-bonded dimer.

### FICD overexpression induces ER stress

We next asked whether FICD-HA overexpression would induce the UPR. To investigate this question in relation to FICD AMPylation activity, we created two mutants of FICD-HA known to increase AMPylation activity by relieving auto-inhibition (E234G) or strongly decrease catalytic activity by replacement of a critical residue in the active site (H363A) [17, 20, 25-27].

To study the potential effects of wild-type or mutant FICD-HA overexpression on the UPR, we analyzed the expression levels of BiP and HERP by qRT-PCR at various time-points after transfection of HEK293 cells (Fig. 5). Treatment of cells with tunicamycin was used as a positive control, and showed a time-dependent increase in BiP and HERP transcript levels. Likewise, overexpression of the constitutively active FICD-HA_E234G_ mutant resulted in increasing levels of BiP and HERP transcripts over time. The same effect, although to a somewhat lesser extent, was observed upon overexpression of FICD-HA_WT_. In contrast, overexpression of the inactive FICD-HA_H363A_ mutant only slightly increased the BiP transcript levels, and did not affect HERP transcripts. Western blot analysis of BiP and HERP confirmed their upregulation at the protein level in a manner closely similar to the transcriptional upregulation, whereas no significant difference in the expression levels of the various FICD-HA constructs was observed (Supplementary S2 Fig.). Based on these results we concluded that UPR induction resulting from FICD-HA overexpression is dependent on and directly linked to the AMPylation activity of the protein, in agreement with similar recent findings [27].

**Fig. 5.**
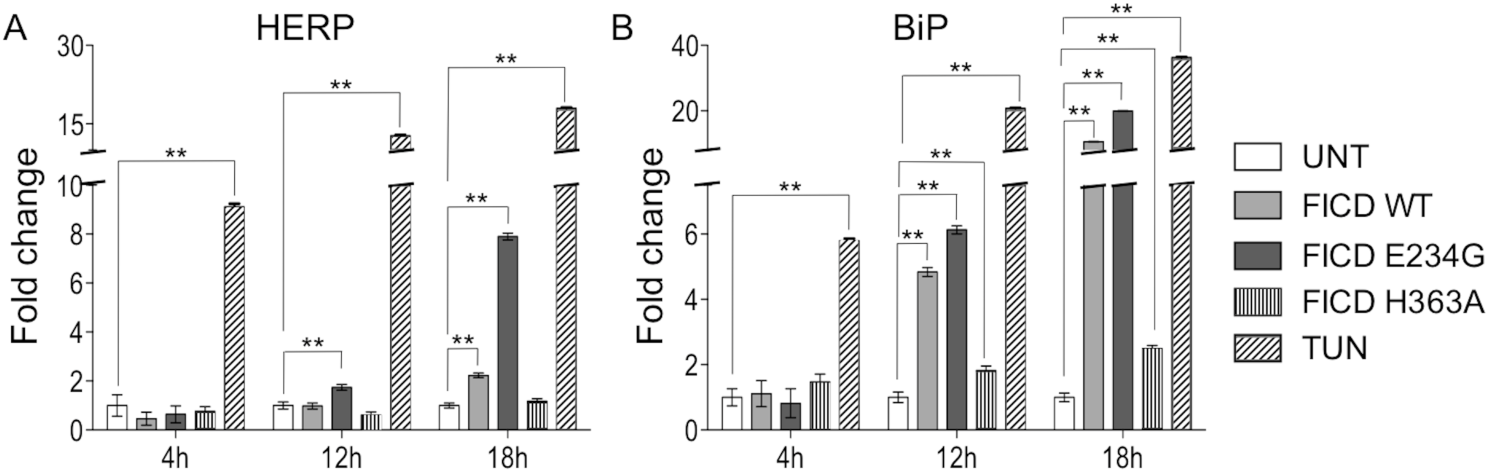
FICD overexpression induces ER stress. (A) Relative abundance of HERP mRNA. Relative gene expression was analyzed by qRT-PCR in HEK293 cells after transfection to express FICD-HA_WT_, FICD-HA_E234G_ or FICD-HA_H363A_. Gene expression was analyzed 4, 12 and 18 h post transfection, respectively. Cells treated with 2.5 μg/ml tunicamycin were used as positive controls. mRNA levels were normalized to actin. Data represent the mean of 3 biological replicates analyzed in technical replicates ± SEM. Statistical analysis was performed using Student’s unpaired *t* test (two tailed, heteroschedastic). (B) Relative abundance of BiP mRNA. The experiment and analysis was performed as in panel (A).

### FICD silencing increases sensitivity to ER stress

To further evaluate the role of FICD in the UPR, we investigated whether siRNA-mediated knockdown of FICD sensitized cells to ER stress induced by a low dose of tunicamycin (2 μg/ml for 24 h) or thapsigargin (5 μM for 24 h). As a positive control, we used knockdown of selenoprotein S/VIMP, which has previously been shown to sensitize cells to treatment with tunicamycin [49]. Sensitivity to ER stress was analyzed with an MTT assay, which measures the metabolic activity of mitochondrial dehydrogenases as readout for cell viability. To knock down FICD, we used two siRNAs designed to target different sequences each providing >80% FICD knock-down after 24 h of transfection as evaluated by qRT-PCR (Supplementary Fig. S1). As indicated by the significant decrease in cell viability (Fig. 6), knockdown of FICD clearly sensitized HEK293 cells to ER stress induced with tunicamycin and thapsigargin providing further evidence for a role of FICD in the UPR. This result is in agreement with previous work that has shown cell death to increase in response to tunicamycin treatment when downregulating FICD [17].

**Fig. 6.**
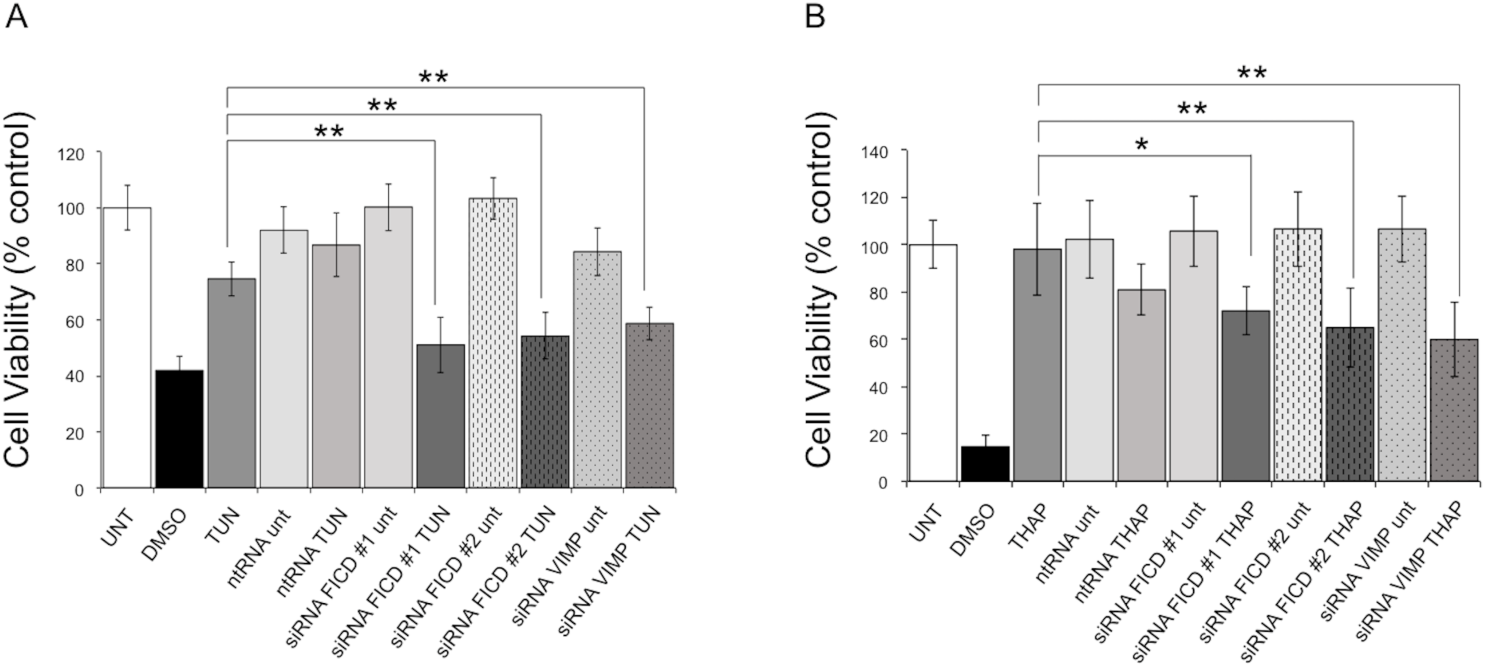
FICD silencing increases sensitivity to ER stress. MTT cell viability assay performed on HEK293 cells treated with the ER stress inducer tunicamycin (A) (TUN; 2 μg/ml for 24 hours) or thapsigargin (B) (THAP; 5μM for 24 hours). Cells treated with 500 mM DMSO were used as positive control for cytotoxicity. RNAi-mediated selenoprotein S/VIMP silencing, known to cause increased sensitivity to ER stress [49], was used for comparison. A non-targeting RNA oligonucleotide (ntRNA) was used as a negative control, and two different oligos (#1 and #2) were used to target FICD. UNT: untreated. Data represent the mean of 6 biological replicates analyzed in 3 technical replicates ± SEM. Statistical analysis was performed using Student’s unpaired *t* test (two tailed, heteroschedastic).

### BiP is the main ER target of AMPylation by FICD

Previous work has identified BiP as a substrate for AMPylation by FICD. Most studies have been performed either *in vitro* using purified components or in whole cell extracts [17, 20, 25, 28]. The cellular identification of BiP as a FICD substrate focused solely on BiP, which precluded information about other potential cellular substrates [27]. Here, we wanted to assess the relative importance of BiP as a substrate for FICD-mediated AMPylation compared to other ER proteins in a native environment.

To identify cellular targets of FICD AMPylation, we isolated the crude membrane fraction containing ER-derived microsomes from cells overexpressing FICD-HA_E234G_ and subjected the intact membranes to an AMPylation assay using [α-^32^P] ATP as the nucleotide source for FICD (as detailed in the Materials and Methods). Protein AMPylation was visualized by autoradiography. Here, we detected one main radioactive band appearing at an apparent molecular weight of ∼75 kDa (Fig. 7A). Subsequent treatment with snake venom type I phosphodiesterase (PDE), which is known to remove AMPylation [50], resulted in a strong decrease in the observed signal. In addition, we detected two weakly labeled proteins at around 45 kDa and 55 kDa, the latter most likely representing FICD auto-AMPylation.

**Fig. 7:**
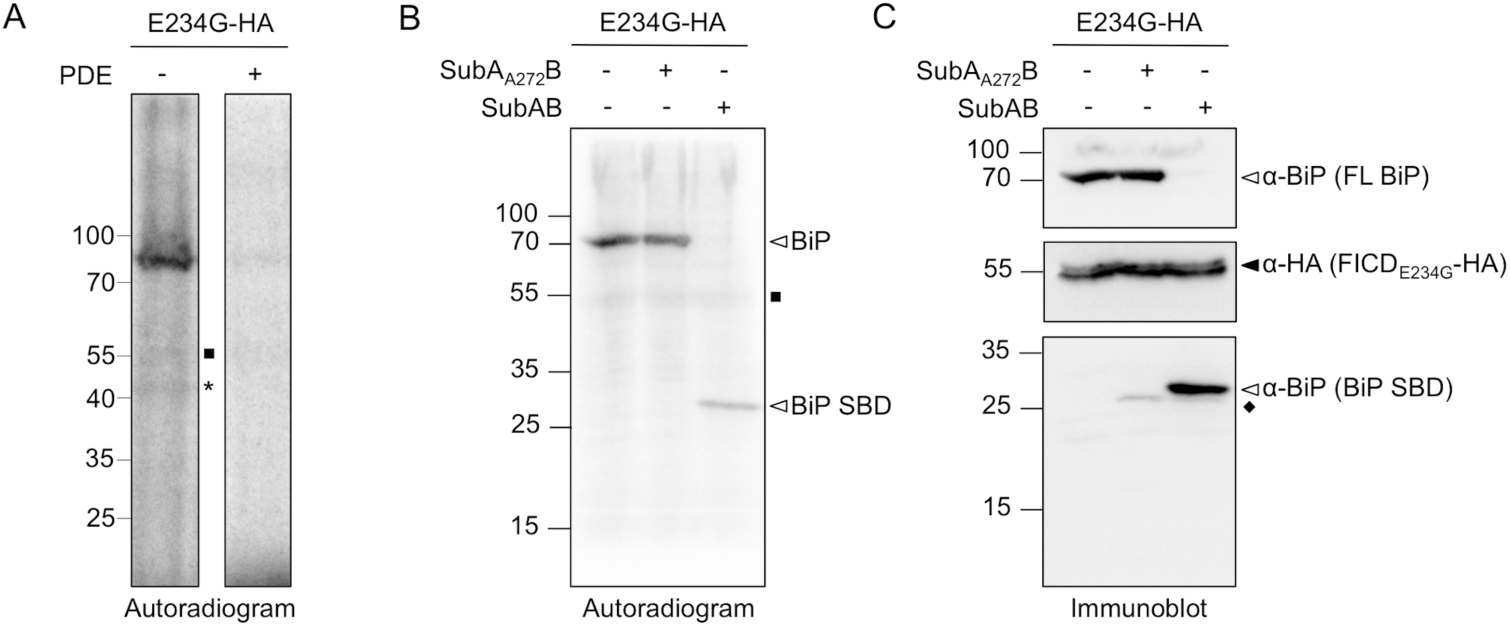
BiP is the main target of FICD AMPylation in ER-derived microsomes. (A) Crude membranes containing ER-derived microsomes prepared from HEK293 cells expressing hyperactive FICD-HA_E234G_ were incubated with [*α*-^32^P]-ATP before solubilization and analysis by non-reducing SDS-PAGE and autoradiography. FICD-HA_E234G_ overexpression gave rise to one main radiolabeled band at around 75 kDa. Upon PDE treatment the AMPylation signal disappeared. Two potential AMPylation targets around 55 kDa (square) and 45 kDa (asterisk) are indicated, the former most likely corresponding to FICD auto-AMPylation. (B) Crude membranes expressing FICD-HA_E234G_ were subjected to the AMPylation assay as above. Solubilized membranes were treated with the subtilase cytotoxin SubAB or the inactive SubA_A272_B mutant. The active SubAB readily cleaved radiolabelled BiP to generate radiolabeled substrate-binding domain (SBD). (C) Immunoblot analysis of the same membrane shown in Panel B probed with an anti-BiP antibody directed against an epitope in the SBD reveals full length BiP (FL BiP) and cleaved SBD levels. A signal from an unknown protein is indicated (diamond).

Given that BiP migrates at 78 kDa, we expected this to be the AMPylated protein detected in Fig. 7A. To confirm BiP as the target of FICD-mediated AMPylation, we transfected cells with FICD-HA_E234G_ and subjected crude membranes to the AMPylation assay described above. After membrane solubilization the lysates were treated with SubAB [51], a subtilase cytotoxin that specifically cleaves BiP at the linker between the nucleotide-binding domain (NBD) and substrate-binding domain (SBD) [52]. The main radioactive band around 75 kDa was readily cleaved by SubAB to generate a radioactively labeled protein migrating at around 28 kDa, and was left uncleaved by the inactive SubA_A272_B mutant (Fig. 7B and C). This finding agrees with recent data where BiP AMPylation was mapped to Thr518 in the SBD of BiP [27], which migrates at 28 kDa. Together our data confirm the identity of the FICD target protein as BiP and suggest that it is the main FICD substrate in the nativelike environment of ER-derived microsomes.

### BiP-HA overexpression increases BiP AMPylation and FICD auto-AMPylation

Next, to investigate the changes in the BiP and FICD AMPylation levels in response to FICD variant and BiP overexpression, we co-transfected cells with plasmids encoding FICD-HA_WT_, FICD-HA_H363A_ or FICD-HA_E234G_, and BiP-HA. As controls we also expressed BiP-HA or FICD-HA_E234G_ alone, or left cells untransfected. We then subjected the crude membranes to an AMPylation assay as described above, followed by immunoprecipitation with an anti-HA antibody. As a reference for IP efficiency and total protein content, 5% of the total microsomal lysate was included on the gels.

In the autoradiogram (Fig. 8A; see also S3 Fig.) we detected AMPylation of both the higher mobility endogenous BiP (5% input lanes) and lower mobility BiP-HA (only visible upon enrichment in the IP lanes). Moreover, hyperactive FICD-HA_E234G_ also displayed (weak) auto-AMPylation (lanes 10 and 12), suggesting that the signal observed in Fig. 7 at around 55 kDa represents FICD auto-AMPylation. Western blots (Fig. 8B and C) show IP efficiency and relative migration of FICD-HA constructs, endogenous BiP and BiP-HA.

**Fig. 8:**
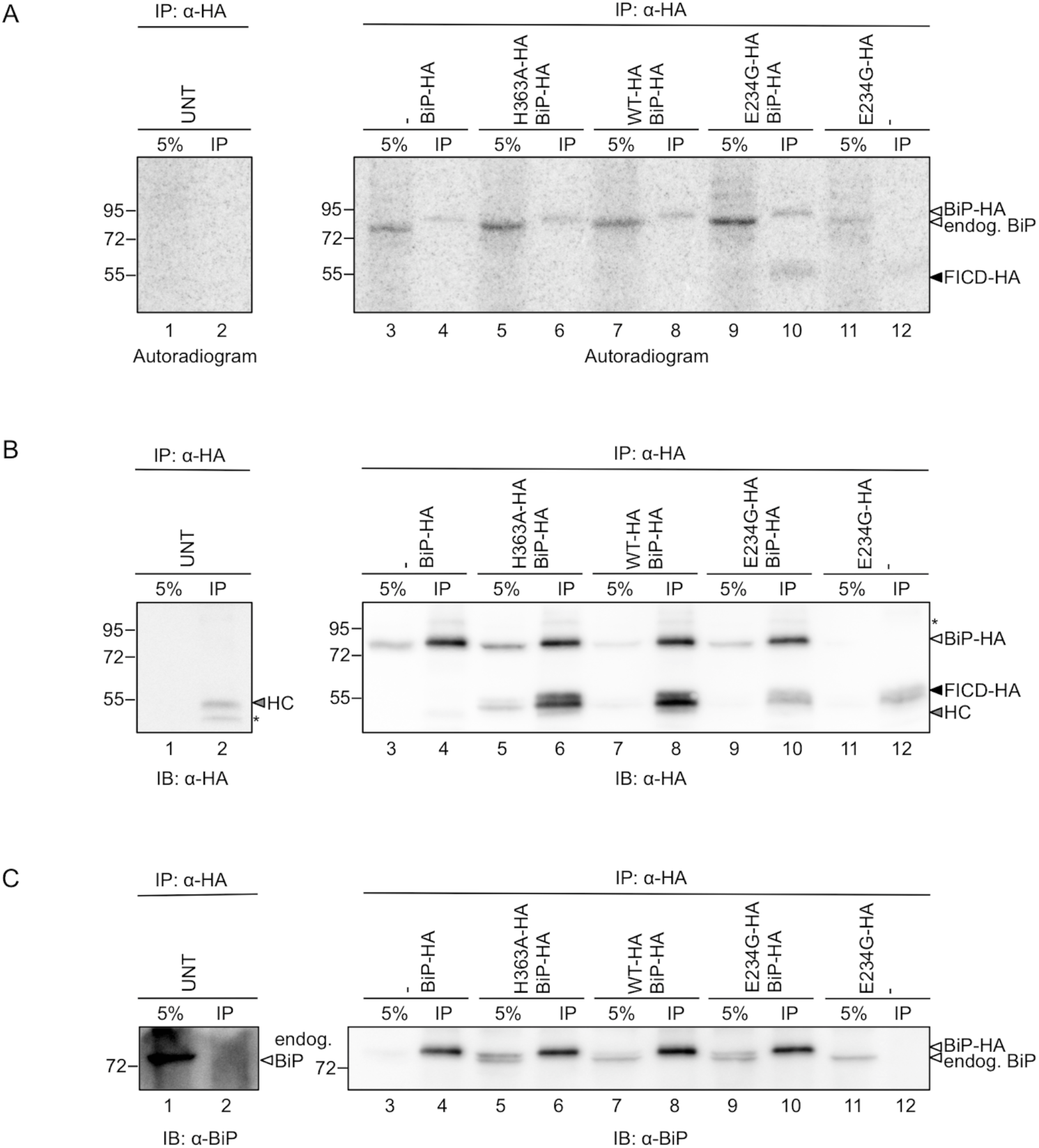
BiP-HA overexpression correlates with increased BiP and FICD AMPylation. HEK293 cells were transfected with combinations of BiP-HA and FICD-HA_WT_, FICD-HA_H363A_ and FICD-HA_E234G_, or left untransfected (UNT), as indicated. Crude membranes were isolated by cell fractionation. The membranes were incubated with [*α*-^32^P]-ATP, chased with an excess of “cold” ATP, EDTA, and EGTA, and solubilised with Triton X-100. 5% of the total lysate was kept as an input reference (5%), and the remaining cleared lysates were subjected to immunoprecipitation by anti-HA antibody (IP). The samples were resolved on a non-reducing SDS-PAGE gel and transferred onto a PVDF membrane for autoradiography (A). The same membrane was blotted with anti-HA (B). Signals from two unknown proteins are labelled with an asterisk. The signal from the antibody heavy chain (HC) is denoted with an arrowhead. The anti-HA blot was directly re-blotted with anti-BiP antibody to detect both endogenous BiP and BiP-HA (C).

Contrary to expectations, we detected BiP AMPylation in the samples containing no exogenous FICD, and when overexpressing inactive (H363A) or FICD-HA_WT_ (lanes 3, 5, 7). In fact, overexpression of BiP-HA alone was sufficient to increase the AMPylation signal of the endogenous BiP (compare lanes 1 and 3). Likewise, co-expression of BiP-HA with FICD-HA_E234G_ (lane 9) gave rise to a higher radioactive signal for endogenous BiP than when FICD-HA_E234G_ was expressed alone (lane 11). Th us, while BiP AMPylation was increased by overexpression of the hyperactive FICD-HA_E234G_ as expected (compare lane 9 with lanes 5 and 7, and lane 11 with lane 1), exogenous expression of BiP-HA resulted in a higher AMPylation of endogenous BiP. In addition to the increase in AMPylation of endogenous BiP, the auto-AMPylation signal of FICD-HA_E234G_ (lanes 10 and 12) was higher when BiP-HA was co-expressed (see also S3 Fig.).

The results in Fig. 7 and Fig. 8 indicated that under the given experimental conditions, BiP is the main ER substrate of FICD. Moreover, BiP-HA overexpression correlated with increased AMPylation of both FICD and BiP itself.

### Abrogation of covalent dimerization increases BiP AMPylation by hyperactive FICD

In a final set of experiments, we studied the effect of cysteine mutations in FICD-HA on AMPylation activity. We expressed FICD-HA_WT_, FICD-HA_C51A/C75A_, FICD-HA_E234G_ and FICD-HA_E234G/C51A/C75A_ in HEK293 cells, and subjected the isolated membranes to the AMPylation assay. Protein levels of endogenous BiP as well as FICD-HA constructs were probed by Western blotting.

While low protein levels of the hyperactive FICD-HA variants (E234G and E234G/C51A/C75A) prevented a direct comparison of BiP AMPylation levels caused by the WT and hyperactive FICD, the analysis showed that mutating both Cys51 and Cys75 to alanine did not impact BiP AMPylation by FICD-HA_WT_ (Fig. 9, lanes 2 and 3; Supplementary S4 Fig.). On the contrary, when mutating both Cys51 and Cys75 to alanine in the context of the hyperactive FICD-HA_E234G_ mutant, BiP AMPylation clearly increased (Fig. 9, lanes 4 and 5; Supplementary S4 Fig.), suggesting that covalent dimerization has the potential to modulate FICD activity.

**Fig. 9:**
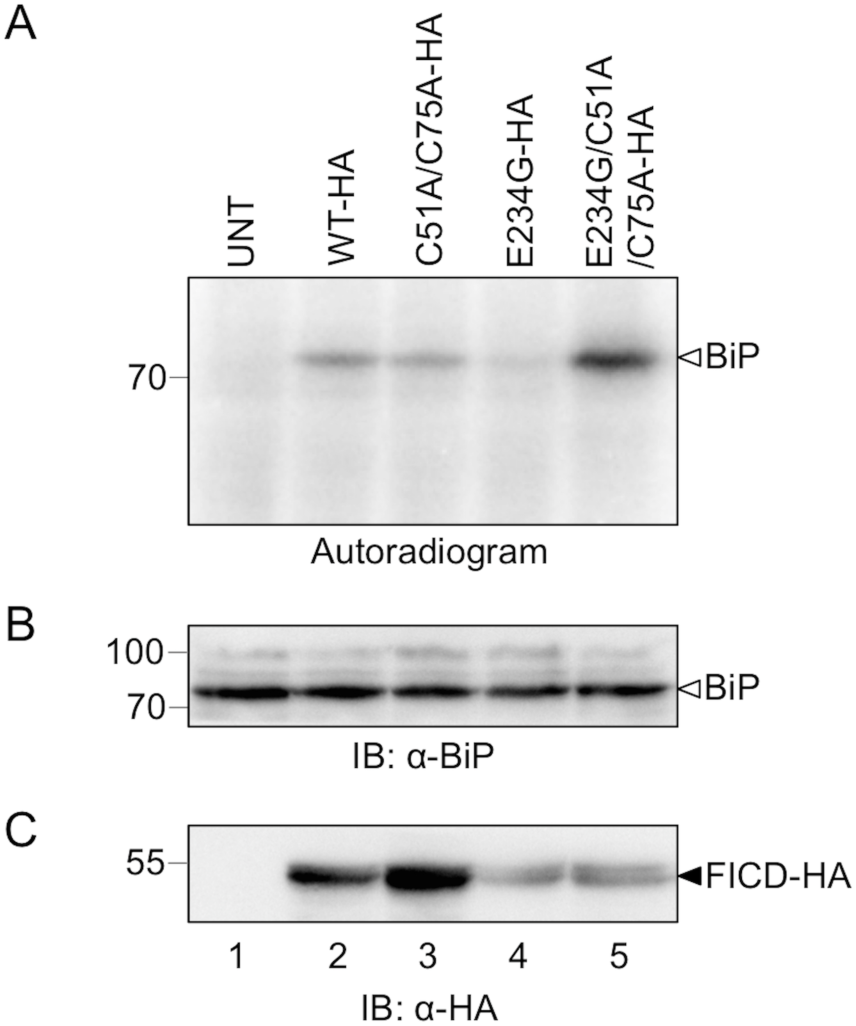
Mutation of Cys51 and Cys75 in FICD-HA_E234G_ increases BiP AMPylation. Crude membranes isolated from untransfected (UNT) cells or HEK293 cells expressing FICD-HA_WT_, FICD-HA_C51A/C75A_, FICD-HA_E234G_ or FICD-HA_E234G/C51A/C75A_ (lanes 1-5), were incubated with [a-^32^P]-ATP before separation by non-reducing SDS-PAGE and autoradiography (A). The membrane was subsequently probed with anti-BiP (B) and anti-HA (to verify the overexpression of the FICD variants; (C). Low protein levels of the hyperactive FICD-HA variants (E234G and E234G/C51A/C75A; see also Supplementary S4 Fig.) precluded a direct comparison of BiP AMPylation levels caused by the “WT enzymes” (FICD-HA or FICD-HA_C51A/C75A_) and the “hyperactive enzymes” (FICD-HA_E234G_ or FICD-HA_E234G/C51A/C75A_). Two independently performed experiments are shown in Supplementary S4 Fig.

## Discussion

Proper regulation of the UPR is of central importance in the cellular decision between stress adaptation and apoptosis, and its misregulation is involved in various diseases such as diabetes, cancer and different neurodegenerative diseases [53-55]. Here, we have performed a cell biological characterization of human FICD, a newly discovered target of the UPR. The finding that human FICD-HA is a type II transmembrane protein of the ER confirms previous findings for *d*FIC obtained in S2 cells [48], where ER localization was inferred from cell fractionation data. The same authors also observed a *d*FIC-HRP fusion protein on the surface of glia cells by electron microscopy [48]. Here, we did not observe any FICD-HA_WT_ on the cell surface of HEK293 cells by confocal microscopy. Rather, in addition to the reticular ER signal, we detected a clear signal at the nuclear rim (Fig. 2, top panel). Likewise, in other recent work endogenous human FICD was shown also to localize to the ER and the nuclear envelope [17, 31]. A similar localization has been demonstrated for the *C. elegans* FIC-1 protein, for which a small fraction was also detected in the cytosol [33]. The physiological consequences of these differences between human FICD, *d*FIC and FIC-1 localization still need to be determined.

We extend our previous finding that FICD expression is upregulated by oxidative ER stress that turns on a mild UPR [16] to show here that FICD is a genuine UPR target. Similar results were published for both *d*FIC [20] and human FICD [17], and have also been found in large-scale studies of the UPR [18, 19]. Moreover, a close functional connection to the UPR is demonstrated by the sensitization of cells to ER stress under conditions of FICD silencing (Fig. 6) [17], as well as the induction of the UPR through overexpression of FICD-HA_WT_ or FICD-HA_E234G_ (Fig. 5). Notably, in contrast to HERP transcript levels that were quite unresponsive to FICD-HA_WT_ overexpression (Fig. 5A), BiP expression levels were readily increased not only by E234G but also by FICD-HA_WT_ overexpression (Fig. 5B). Given the ability of FICD to de-AMPylate BiP at low ER protein load [37], we speculate that increased levels of FICD could activate BiP, which would induce ER stress and thus increase BiP transcript levels as observed. Overexpression of FICD E234G has previously been demonstrated to negatively affect cell viability and induce cell death [17, 27, 31, 37].

The mechanism underlying these observations likely relates to FICD-mediated inactivation of BiP by AMPylation [20, 27] (although it has also been reported that AMPylation activates BiP ATPase activity [17]). Thus, when inactivated by AMPylation through overexpression of FICD E234G, the cell responds by activating the UPR to create a pool of active BiP [27]. Eventually, the cell can no longer cope with unchecked FICD AMPylation activity and undergoes apoptosis. Moreover, FICD knockout has been shown to cause a delay in the activation of the UPR sensors PERK and IRE1[27], as a result of excess active BiP. Whether AMPylation directly modulates the interaction of BiP with the UPR sensor proteins in addition to interfering with BiP chaperone activity to influence UPR activation remains to be seen.

It is thus clear that the cell closely monitors the level of AMPylation-active FICD, which in turn influences BiP levels and activity to modulate the UPR. Here we show (Fig. 8, S3 Fig.) that increasing the cellular pool of BiP by exogenous expression causes two effects: i) more endogenous BiP becomes AMPylated and ii) FICD-HA_E234G_ auto-AMPylation increases. The latter result suggests that the observed increase in BiP AMPylation resulting from BiP-HA overexpression is not merely due to a potential increase in the cellular pool of endogenous BiP, but likely associated with an increase in FICD AMPylation activity. It is noteworthy that the increase in BiP AMPylation occurs independently of FICD overexpression, indicating that the activity of endogenous FICD can be regulated solely by the cellular levels of its substrate BiP. This principle apparently also applies to the level of FICD auto-AMPylation (at least for the hyperactive E234G mutant). Whether auto-AMPylation represents a side effect of FICD activity, or actively partakes in its regulation remains to be seen. Overall, we propose that excess BiP is inactivated by AMPylation in order to maintain an optimal level of the active protein that allows it to carry out its functions in protein folding as well as UPR signaling.

Despite the recent advances in understanding the cellular function of FICD in relation to BiP, it has been unclear whether other significant cellular substrates exist. For instance, a number of potential substrates were identified in a whole cell extract upon AMPylation with purified recombinant FICD, but no other ER proteins than BiP were identified [28]. Here, we demonstrate that in the native-like setting of ER-derived microsomes isolated from HEK293 cells, BiP is clearly the predominant target of FICD-HA AMPylation. While FICD potentially has additional substrates of low abundance, AMPylated below the detection level of our assay (with a band at ∼45 kDa as a possible exception; Fig. 7), the present data argue that BiP is indeed the main ER substrate of FICD and suggest that the primary biological function of FICD, at least in this cellular system, is to control the AMPylation status of BiP. The same seems to be the case in S2 cells, where at steady state BiP is the predominant protein recognized by an antibody raised against AMPylated threonine [20].

The BiP AMPylation site has been a matter of some debate, with residues identified as target sites both in the NBD and SBD, depending on the experimental system [17, 20, 27]. In the present work, we observed BiP AMPylation on the SBD of BiP (Fig. 7), in agreement with the cell biological characterization performed by the Ron lab [27].

Our work further demonstrates that FICD-HA forms disulfide-bonded dimers when expressed in HEK293 cells. Unfortunately, we have not yet been able to identify a commercial FICD antibody that convincingly detects the endogenous protein by Western blotting. Neither have our efforts to use affinity purified anti-FICD serum for detection of endogenous FICD been successful. While we can therefore not exclude that the observed dimer appears as a result of FICD overexpression, we routinely observe a large fraction (about half) of the protein in the dimeric form as judged by Western blotting (see e.g. Fig. 4). Moreover, we see little evidence of other disulfide-bonded species involving FICD-HA. Based on these observations, our findings likely reflect the situation for the endogenous protein.

The crystal structure of human FICD shows that the protein forms a non-covalent dimer, not only in the crystals but also in solution [25]. FIC-1 also crystallizes as a non-covalent dimer [33]. The structural organization of the dimer is such that the interface is constituted exclusively by residues from the Fic domain, and with the two TPR regions placed at either end of the dimer pointing away from each other [25]. Although the evolutionarily strictly conserved Cys421 is positioned relatively close to the dimer interface, it does not engage in a disulfide bond in the crystal structure, a finding that was confirmed here in human cells (Fig. 4). Instead, we find by MS analysis that in HEK293 cells FICD-HA_WT_ Cys75 forms a disulfide with Cys75 in a neighboring molecule of FICD-HA_WT_. Based on the finding that only the FICD-HA_C51A/C75A_ double mutant did not form disulfide-linked dimers, while the FICD-HA_C51A_ and FICD-HA_C75A_ single cysteine mutants were still able to do so, Cys51-Cys51 disulfide-bonds likely also form. In terms of evolutionary conservation, Cys51 is very well conserved among species, whereas Cys75 is less conserved. Only very few species are apparently missing both cysteines in the protein; these proteins then often harbor a Cys in a proximal position that potentially fulfills the same function as Cys51 and Cys75. The most notable exception seems to be the distantly related *d*FIC, which does not harbor any cysteines in this region of the protein.

Disrupting covalent dimerization is apparently not sufficient to turn on the AMPylation activity of the wild-type enzyme. Intriguingly, FICD-HA_E234G/C51A/C75A_ on the other hand displays higher AMPylation activity towards BiP than FICD-HA_E234G_ (Fig. 8), suggesting that preventing covalent dimerization either directly increases AMPylation activity or allows binding of a partner protein that promotes AMPylation by FICD E234G. Whether FICD activity is redox regulated, and thus potentially modulated by the action of PDIs as is the case for the proximal UPR sensors ATF6 and IRE1 [56], remains to be seen.

## Author Contributions

Conceived and designed the experiments: RM MSV CMH CS BC JJE ESS LE. Performed the experiments: RM MSV CMH CLS APC YL CS BC. Analyzed the data: RM MSV CMH CS JJE BC ESS LE. Contributed reagents: JP AWP. Wrote the paper: RM MSV CMH LE. Edited the manuscript: RM MSV CMH CLS APC YL CS JP AWP ESS LE.

## Acknowledgments

We thank Linda Hendershot and Ari Helenius for providing antibodies, and Alexander Harms, Laura Cesa and all members of the Ellgaard lab for critical reading of the manuscript.

---

**S1 Fig.**
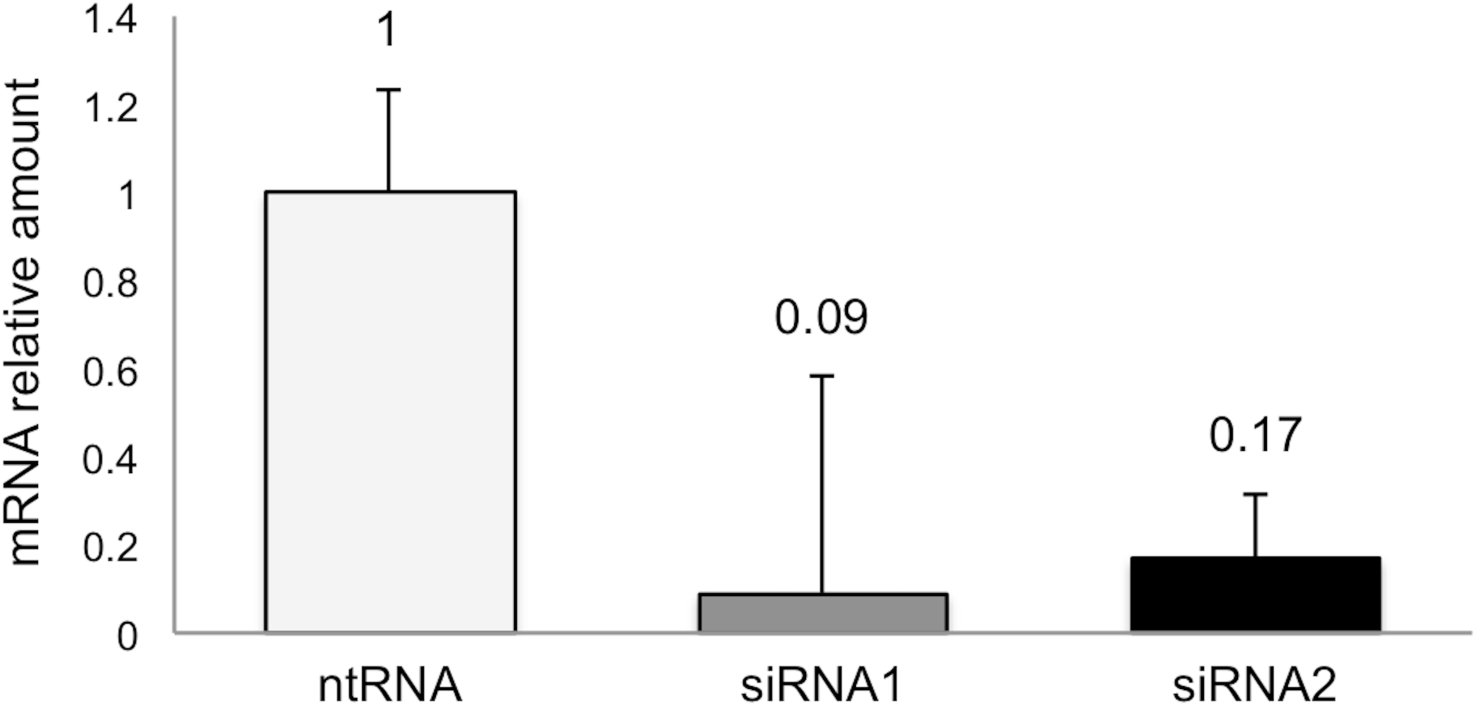
Relative abundance of FICD mRNA after siRNA-mediated silencing. Relative abundance of FICD mRNA in HEK293 cells analyzed by qRT-PCR. Cells were treated for 48 h with two different siRNAs targeting FICD (Hs_FICD_1 and Hs_FICD_2) at a concentration of 5 nM. mRNA levels were normalized to actin. As negative control, cells were transfected with 10 nM non-targeting siRNA (ntRNA). Data represent the mean of 2 biological replicates analyzed in 3 technical replicates ± SEM. Numbers indicate the remaining fraction of FICD mRNA.

**S2 Fig.**
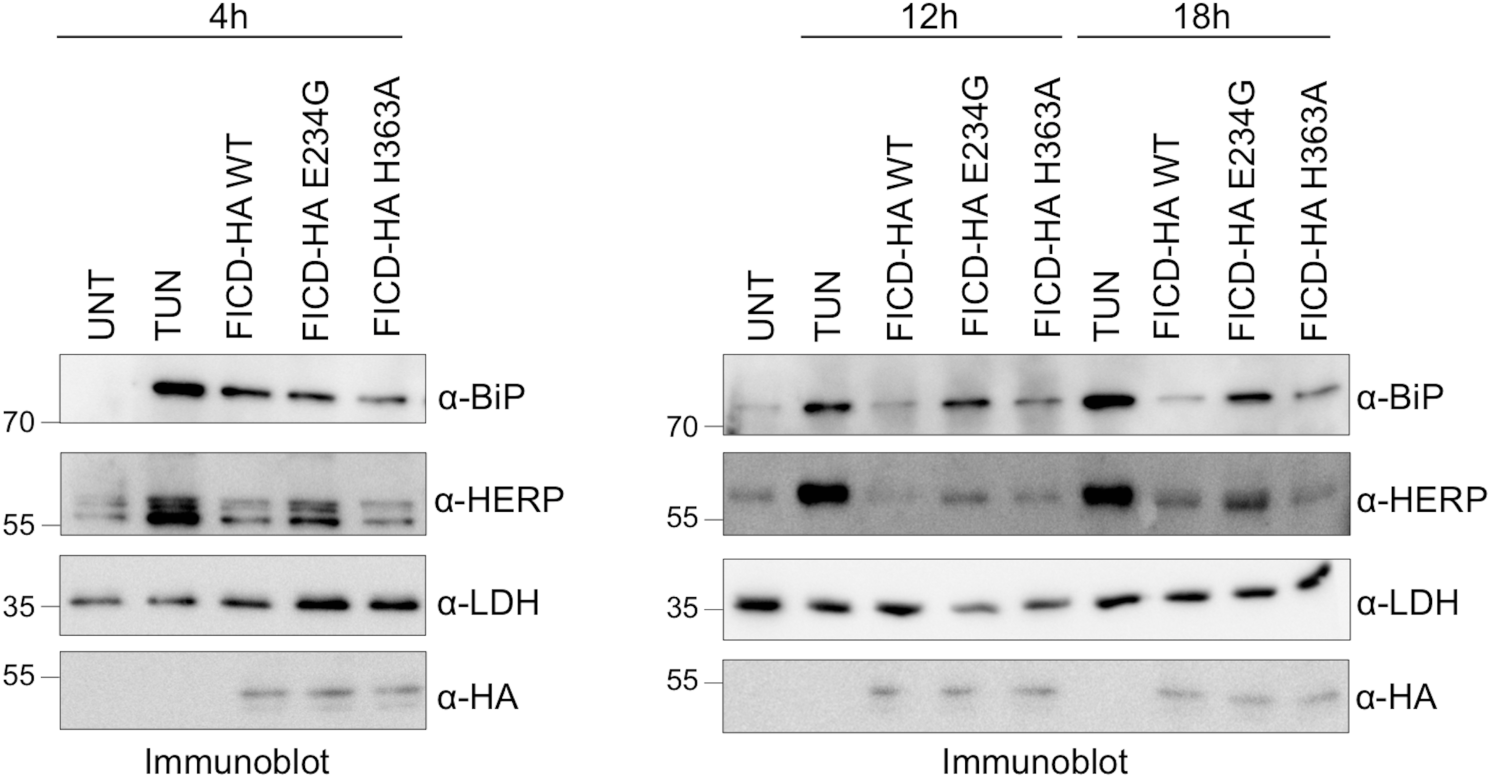
FICD overexpression induces ER stress. Cells expressing FICD-HA_WT_, FICD-HA_E234G_ and FICD-HA_H363A_ as indicated, were analyzed by Western blotting at different time points after transfection to determine the protein levels of BiP and HERP. LDH was used as loading control. Cells treated with 2.5 μg/ml tunicamycin for 8 hours were used to generate positive control lysates for UPR induction. Anti-HA immunoblotting shows the level of transfection.

**S3 Fig.**
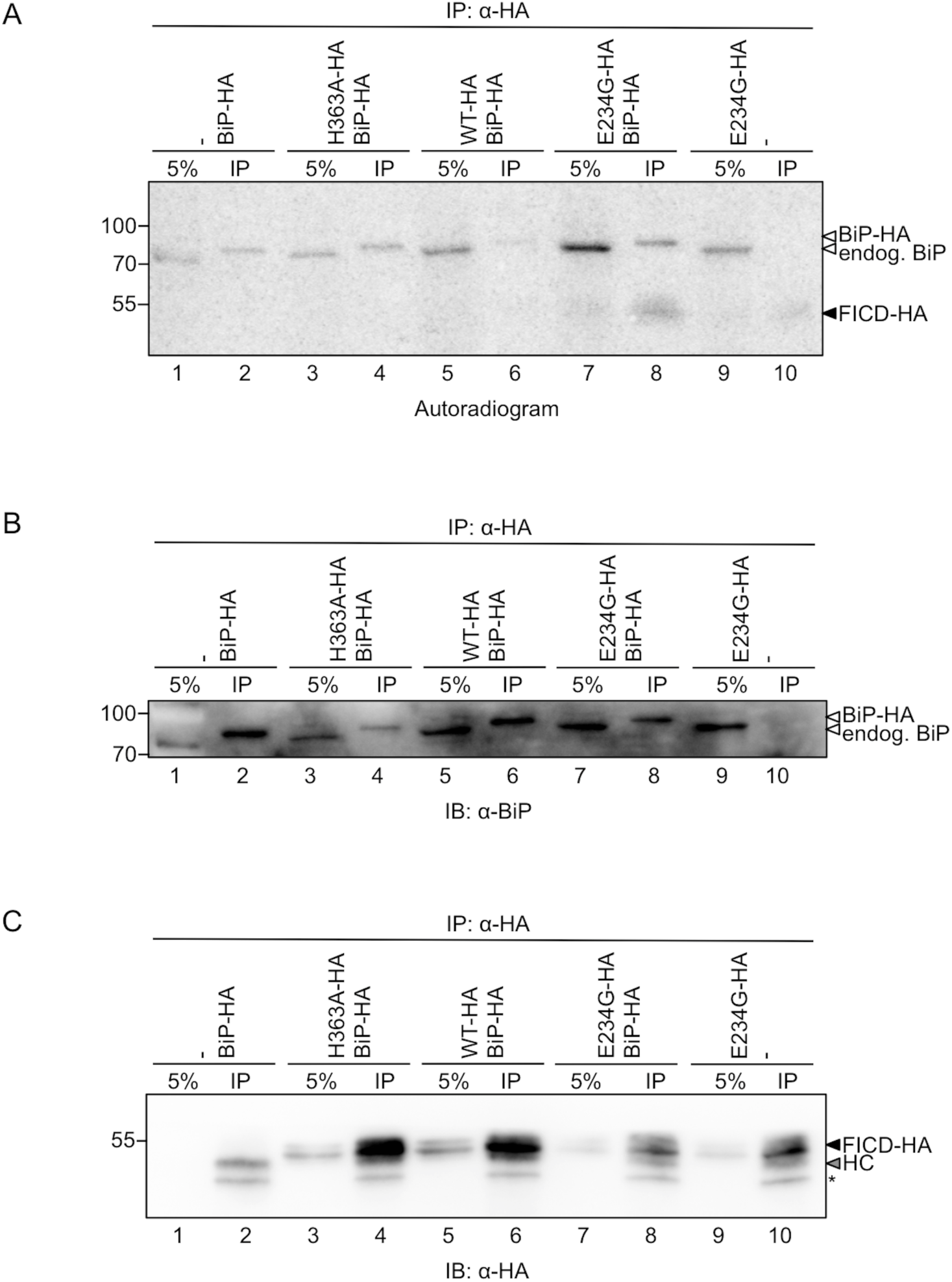
BiP-HA overexpression correlates with increased BiP and FICD AMPylation. HEK293 cells were transfected with combinations of BiP-HA and FICD-HA_WT_, FICD-HA_H363A_ and FICD-HA_E234G_, as indicated. Crude membranes were isolated by cell fractionation. The membranes were incubated with [*α*-^32^P]-ATP, chased with an excess of “cold” ATP, EDTA, and EGTA, and solubilised with Triton X-100. 5% of the total lysate was kept as an input reference (5%), and the remaining cleared lysates were subjected to immunoprecipitation by anti-HA antibody (IP). The samples were resolved on a non-reducing SDS-PAGE gel and transferred onto a PVDF membrane for autoradiography (A). The same membrane was blotted with anti-BiP (B) and anti-HA (C). Signal from an unknown protein is labelled with an asterisk. The signal from the antibody heavy chain (HC) is denoted with an arrowhead.

**S4 Fig:**
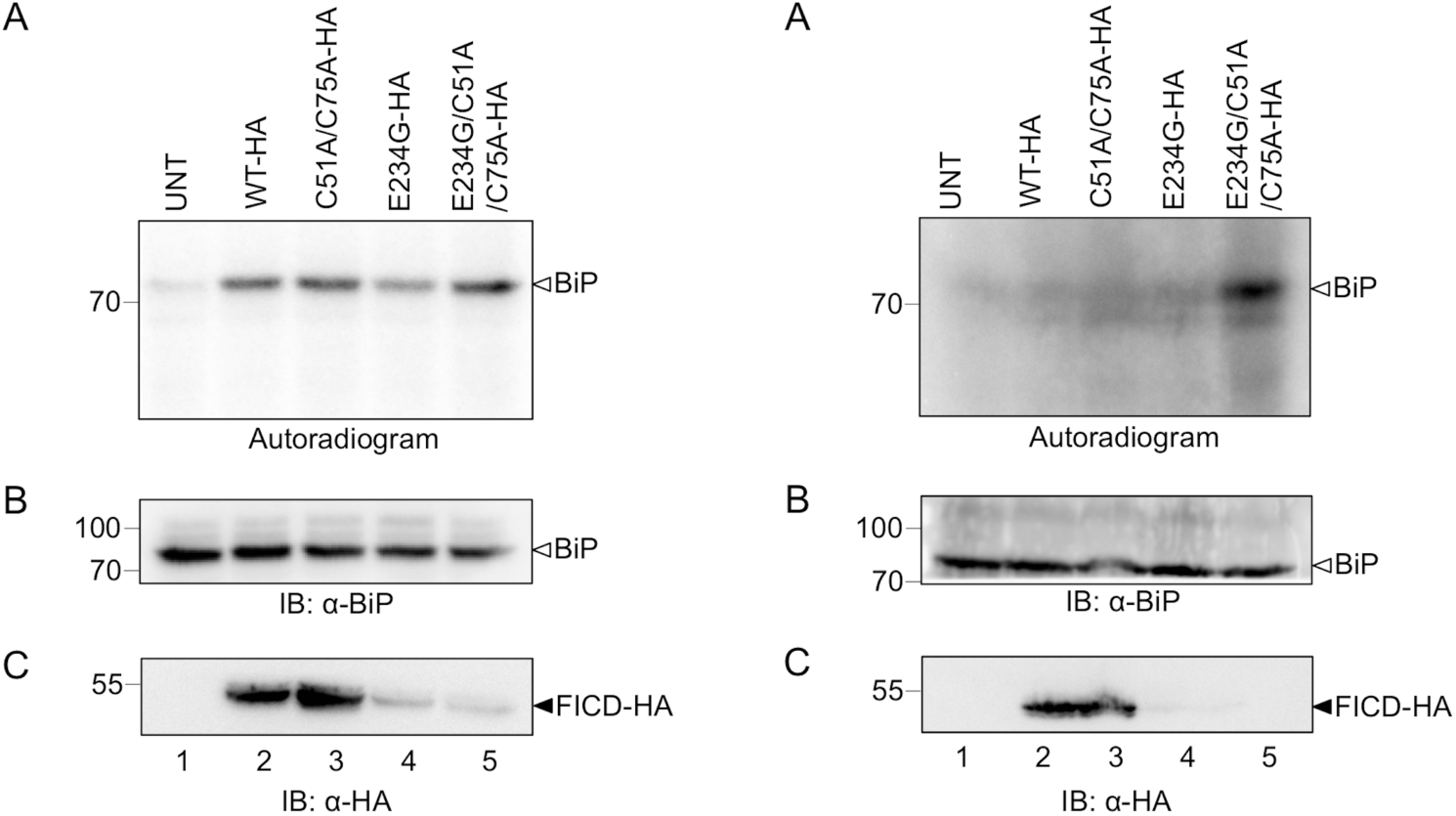
Mutation of Cys51 and Cys75 in FICD-HA_E234G_ increases BiP AMPylation. Crude membranes isolated from untransfected (UNT) cells or HEK293 cells expressing FICD-HA_WT_, FICD-HA_C51A/C75A_, FICD-HA_E234G_ or FICD-HA_E234G/C51A/C75A_ (lanes 1-5), were incubated with [*α*-^32^P]-ATP before separation by non-reducing SDS-PAGE and autoradiography (A). The membrane was subsequently probed with anti-BiP (B) and anti-HA (to verify the overexpression of the FICD variants; (C). Low protein levels of the hyperactive FICD-HA variants (E234G and E234G/C51A/C75A) precluded a direct comparison of BiP AMPylation levels caused by the “WT enzymes” (FICD-HA or FICD-HA_C51A/C75A_) and the “hyperactive enzymes” (FICD-HA_E234G_ or FICD-HA_E234G/C51A/C75A_). Two independently performed experiments are shown.

## References

1. Gardner BM, Pincus D, Gotthardt K, Gallagher CM, Walter P. Endoplasmic reticulum stress sensing in the unfolded protein response. Cold Spring Harb Perspect Biol. 2013; 5(3): a013169. doi: 10.1101/cshperspect.a013169. PubMed PMID: 23388626.

2. Volmer R, Ron D. Lipid-dependent regulation of the unfolded protein response. Curr Opin Cell Biol. 2014;33C:67–73. doi: 10.1016/j.ceb.2014.12.002. PubMed PMID: 25543896.

3. Harding HP, Zhang Y, Ron D. Protein translation and folding are coupled by an endoplasmic-reticulum-resident kinase. <u>Nature</u>. 1999;397(6716):271–4. doi: 10.1038/16729. PubMed PMID: 9930704.

4. Hollien J, Weissman JS. Decay of endoplasmic reticulum-localized mRNAs during the unfolded protein response. Science. 2006;313(5783):104–7. doi:10.1126/science.1129631. PubMed PMID: 16825573.

5. Cox JS, Walter P. A novel mechanism for regulating activity of a transcription factor that controls the unfolded protein response. Cell. 1996; 87(3): 391–404. PubMed PMID: 8898193.

6. Yoshida H, Matsui T, Yamamoto A, Okada T, Mori K. XBP1 mRNA is induced by ATF6 and spliced by IRE1 in response to ER stress to produce a highly active transcription factor. Cell. 2001; 107(7): 881–91. PubMed PMID: 11779464.

7. Haze K, Yoshida H, Yanagi H, Yura T, Mori K. Mammalian transcription factor ATF6 is synthesized as a transmembrane protein and activated by proteolysis in response to endoplasmic reticulum stress. Mol Biol Cell. 1999; 10(11): 3787–99. PubMed PMID: 10564271; PubMed Central PMCID: PMC25679.

8. Otero JH, Lizak B, Hendershot LM. Life and death of a BiP substrate. Semin Cell Dev Biol. 2010; 21(5): 472–8. doi: 10.1016/j.semcdb.2009.12.008. PubMed PMID: 20026282; PubMed Central PMCID: PMC2883687.

9. Ohta M, Takaiwa F. Emerging features of ER resident J-proteins in plants. Plant Signal Behav. 2014; 9(7): e28194. doi: 10.4161/psb.28194. PubMed PMID: 25763480.

10. Jin Y, Awad W, Petrova K, Hendershot LM. Regulated release of ERdj3 from unfolded proteins by BiP. EMBO J. 2008; 27(21): 2873–82. doi: 10.1038/emboj.2008.207. PubMed PMID: 18923428; PubMed Central PMCID: PMC2580786.

11. Shen Y, Meunier L, Hendershot LM. Identification and characterization of a novel endoplasmic reticulum (ER) DnaJ homologue, which stimulates ATPase activity of BiP in vitro and is induced by ER stress. J Biol Chem. 2002; 277(18): 15947–56. doi: 10.1074/jbc.M112214200. PubMed PMID: 11836248.

12. Cunnea PM, Miranda-Vizuete A, Bertoli G, Simmen T, Damdimopoulos AE, Hermann S, et al. ERdj5, an endoplasmic reticulum (ER)-resident protein containing DnaJ and thioredoxin domains, is expressed in secretory cells or following ER stress. J Biol Chem. 2003; 278(2): 1059–66. doi: 10.1074/jbc.M206995200. PubMed PMID: 12411443.

13. Hosoda A, Kimata Y, Tsuru A, Kohno K. JPDI, a novel endoplasmic reticulum-resident protein containing both a BiP-interacting J-domain and thioredoxin-like motifs. J Biol Chem. 2003; 278(4): 2669–76. doi: 10.1074/jbc.M208346200. PubMed PMID: 12446677.

14. Dufey E, Sepulveda D, Rojas-Rivera D, Hetz C. Cellular mechanisms of endoplasmic reticulum stress signaling in health and disease. 1. An overview. Am J Physiol Cell Physiol. 2014;307(7):C582–94. doi: 10.1152/ajpcell.00258.2014. PubMed PMID: 25143348.

15. Carrara M, Prischi F, Nowak PR, Kopp MC, Ali MM. Noncanonical binding of BiP ATPase domain to Ire1 and Perk is dissociated by unfolded protein CH1 to initiate ER stress signaling. eLife. 2015;4. doi: 10.7554/eLife.03522. PubMed PMID: 25692299; PubMed Central PMCID: PMC4337721.

16. Hansen HG, Schmidt JD, Soltoft CL, Ramming T, Geertz-Hansen HM, Christensen B, et al. Hyperactivity of the Ero1alpha oxidase elicits endoplasmic reticulum stress but no broad antioxidant response. J Biol Chem. 2012; 287(47): 39513–23. doi: 10.1074/jbc.M112.405050. PubMed PMID: 23027870; PubMed Central PMCID: PMC3501080.

17. Sanyal A, Chen AJ, Nakayasu ES, Lazar CS, Zbornik EA, Worby CA, et al. A Novel Link Between Fic (Filamentation induced by cAMP)-mediated Adenylylation/AMPylation and the Unfolded Protein Response. J Biol Chem. 2015. doi: 10.1074/jbc.M114.618348. PubMed PMID: 25601083.

18. Shoulders MD, Ryno LM, Genereux JC, Moresco JJ, Tu PG, Wu C, et al. Stress-independent activation of XBP1s and/or ATF6 reveals three functionally diverse ER proteostasis environments. Cell Rep. 2013; 3(4): 1279–92. doi: 10.1016/j.celrep.2013.03.024. PubMed PMID: 23583182; PubMed Central PMCID: PMC3754422.

19. Dombroski BA, Nayak RR, Ewens KG, Ankener W, Cheung VG, Spielman RS. Gene expression and genetic variation in response to endoplasmic reticulum stress in human cells. Am J Hum Genet. 2010; 86(5): 719–29. doi: 10.1016/j.ajhg.2010.03.017. PubMed PMID: 20398888; PubMed Central PMCID: PMC2869002.

20. Ham H, Woolery AR, Tracy C, Stenesen D, Kramer H, Orth K. Unfolded protein response-regulated dFic reversibly AMPylates BiP during endoplasmic reticulum homeostasis. The Journal of biological chemistry. 2014. doi: 10.1074/jbc.M114.612515. PubMed PMID: 25395623.

21. Garcia-Pino A, Zenkin N, Loris R. The many faces of Fic: structural and functional aspects of Fic enzymes. Trends Biochem Sci. 2014; 39(3): 121–9. doi: 10.1016/j.tibs.2014.01.001. PubMed PMID: 24507752.

22. Harms A, Maisonneuve E, Gerdes K. Mechanisms of bacterial persistence during stress and antibiotic exposure. Science. 2016;354(6318). doi: 10.1126/science.aaf4268. PubMed PMID: 27980159.

23. Woolery AR, Luong P, Broberg CA, Orth K. AMPylation: Something Old is New Again. Front Microbiol. 2010; 1: 1–6. doi: 10.3389/fmicb.2010.00113. PubMed PMID: 21607083; PubMed Central PMCID: PMC3095399.

24. Zeytuni N, Zarivach R. Structural and functional discussion of the tetra-trico-peptide repeat, a protein interaction module. Structure. 2012; 20(3): 397–405. doi: 10.1016/j.str.2012.01.006. PubMed PMID: 22404999.

25. Bunney TD, Cole AR, Broncel M, Esposito D, Tate EW, Katan M. Crystal structure of the human, FIC-domain containing protein HYPE and implications for its functions. Structure. 2014; 22(12): 1831–43. doi: 10.1016/j.str.2014.10.007. PubMed PMID: 25435325; PubMed Central PMCID: PMC4342408.

26. Engel P, Goepfert A, Stanger FV, Harms A, Schmidt A, Schirmer T, et al. Adenylylation control by intra- or intermolecular active-site obstruction in Fic proteins. Nature. 2012;482(7383):107–10. doi: 10.1038/nature10729. PubMed PMID: 22266942.

27. Preissler S, Rato C, Chen R, Antrobus R, Ding S, Fearnley IM, et al. AMPylation matches BiP activity to client protein load in the endoplasmic reticulum. eLife. 2015;4. doi: 10.7554/eLife.12621. PubMed PMID: 26673894.

28. Broncel M, Serwa RA, Bunney TD, Katan M, Tate EW. Global profiling of HYPE mediated AMPylation through a chemical proteomic approach. Mol Cell Proteomics. 2015. doi: 10.1074/mcp.O115.054429. PubMed PMID: 26604261.

29. Yu X, LaBaer J. High-throughput identification of proteins with AMPylation using self-assembled human protein (NAPPA) microarrays. Nat Protoc. 2015; 10(5): 756–67. doi: 10.1038/nprot.2015.044. PubMed PMID: 25881200; PubMed Central PMCID: PMCPMC4464792.

30. Truttmann MC, Zheng X, Hanke L, Damon JR, Grootveld M, Krakowiak J, et al. Unrestrained AMPylation targets cytosolic chaperones and activates the heat shock response. Proc Natl Acad Sci USA. 2017;114(2):E152–E60. doi: 10.1073/pnas.1619234114. PubMed PMID: 28031489; PubMed Central PMCID: PMCPMC5240723.

31. Truttmann MC, Wu Q, Stiegeler S, Duarte JN, Ingram J, Ploegh HL. HypE-specific Nanobodies as Tools to modulate HypE-mediated Target AMPylation. J Biol Chem. 2015. doi: 10.1074/jbc.M114.634287. PubMed PMID: 25678711.

32. Lewallen DM, Sreelatha A, Dharmarajan V, Madoux F, Chase P, Griffin PR, et al. Inhibiting AMPylation: a novel screen to identify the first small molecule inhibitors of protein AMPylation. ACS Chem Biol. 2014; 9(2): 433–42. doi: 10.1021/cb4006886. PubMed PMID: 24274060; PubMed Central PMCID: PMC3944102.

33. Truttmann MC, Cruz VE, Guo X, Engert C, Schwartz TU, Ploegh HL. The Caenorhabditis elegans Protein FIC-1 Is an AMPylase That Covalently Modifies Heat-Shock 70 Family Proteins, Translation Elongation Factors and Histones. PLoS Genet. 2016; 12(5): e1006023. doi: 10.1371/journal.pgen.1006023. PubMed PMID: 27138431; PubMed Central PMCID: PMCPMC4854385.

34. Chambers JE, Petrova K, Tomba G, Vendruscolo M, Ron D. ADP ribosylation adapts an ER chaperone response to short-term fluctuations in unfolded protein load. J Cell Biol. 2012; 198(3): 371–85. doi: 10.1083/jcb.201202005. PubMed PMID: 22869598; PubMed Central PMCID: PMC3413365.

35. Carlsson L, Lazarides E. ADP-ribosylation of the Mr 83,000 stress-inducible and glucose-regulated protein in avian and mammalian cells: modulation by heat shock and glucose starvation. Proc Natl Acad Sci USA. 1983; 80(15): 4664–8. PubMed PMID: 6576354; PubMed Central PMCID: PMC384104.

36. Hendershot LM, Ting J, Lee AS. Identity of the immunoglobulin heavy-chain-binding protein with the 78,000-dalton glucose-regulated protein and the role of posttranslational modifications in its binding function. Mol Cell Biol. 1988; 8(10): 4250–6. PubMed PMID: 3141786; PubMed Central PMCID: PMC365497.

37. Preissler S, Rato C, Perera LA, Saudek V, Ron D. FICD acts bifunctionally to AMPylate and de-AMPylate the endoplasmic reticulum chaperone BiP. Nat Struct Mol Biol. 2016. doi: 10.1038/nsmb.3337. PubMed PMID: 27918543.

38. Shen J, Chen X, Hendershot L, Prywes R. ER stress regulation of ATF6 localization by dissociation of BiP/GRP78 binding and unmasking of Golgi localization signals. Dev Cell. 2002; 3(1): 99–111. PubMed PMID: 12110171.

39. Appenzeller-Herzog C, Ellgaard L. In vivo reduction-oxidation state of protein disulfide isomerase: the two active sites independently occur in the reduced and oxidized forms. Antioxid Redox Signal. 2008; 10(1): 55–64. doi: 10.1089/ars.2007.1837. PubMed PMID: 17939758.

40. Dreszer TR, Karolchik D, Zweig AS, Hinrichs AS, Raney BJ, Kuhn RM, et al. The UCSC Genome Browser database: extensions and updates 2011. Nucleic Acids Res. 2012;40(Database issue):D918–23. doi: 10.1093/nar/gkr1055. PubMed PMID: 22086951; PubMed Central PMCID: PMC3245018.

41. Rambaldi D, Ciccarelli FD. FancyGene: dynamic visualization of gene structures and protein domain architectures on genomic loci. Bioinformatics. 2009; 25(17): 2281–2. doi: 10.1093/bioinformatics/btp381. PubMed PMID: 19542150; PubMed Central PMCID: PMC2734320.

42. Haugstetter J, Blicher T, Ellgaard L. Identification and characterization of a novel thioredoxin-related transmembrane protein of the endoplasmic reticulum. J Biol Chem. 2005; 280(9): 8371–80. doi: 10.1074/jbc.M413924200. PubMed PMID: 15623505.

43. Shevchenko A, Tomas H, Havlis J, Olsen JV, Mann M. In-gel digestion for mass spectrometric characterization of proteins and proteomes. Nat Protoc. 2006; 1(6): 2856–60. doi: 10.1038/nprot.2006.468. PubMed PMID: 17406544.

44. Rappsilber J, Ishihama Y, Mann M. Stop and go extraction tips for matrix-assisted laser desorption/ionization, nanoelectrospray, and LC/MS sample pretreatment in proteomics. Anal Chem. 2003; 75(3): 663–70. PubMed PMID: 12585499.

45. Djebali S, Davis CA, Merkel A, Dobin A, Lassmann T, Mortazavi A, et al. Landscape of transcription in human cells. Nature. 2012;489(7414):101–8. doi: 10.1038/nature11233. PubMed PMID: 22955620; PubMed Central PMCID: PMC3684276.

46. Yoshida H, Haze K, Yanagi H, Yura T, Mori K. Identification of the cis-acting endoplasmic reticulum stress response element responsible for transcriptional induction of mammalian glucose-regulated proteins. Involvement of basic leucine zipper transcription factors. J Biol Chem. 1998; 273(50): 33741–9. PubMed PMID: 9837962.

47. Yamamoto K, Sato T, Matsui T, Sato M, Okada T, Yoshida H, et al. Transcriptional induction of mammalian ER quality control proteins is mediated by single or combined action of ATF6alpha and XBP1. Developmental cell. 2007; 13(3): 365–76. doi: 10.1016/j.devcel.2007.07.018. PubMed PMID: 17765680.

48. Rahman M, Ham H, Liu X, Sugiura Y, Orth K, Kramer H. Visual neurotransmission in Drosophila requires expression of Fic in glial capitate projections. Nature neuroscience. 2012; 15(6): 871–5. doi: 10.1038/nn.3102. PubMed PMID: 22544313; PubMed Central PMCID: PMC3578554.

49. Lee JH, Kwon JH, Jeon YH, Ko KY, Lee SR, Kim IY. Pro178 and Pro183 of selenoprotein S are essential residues for interaction with p97(VCP) during endoplasmic reticulum-associated degradation. The Journal of biological chemistry. 2014;289(20): 13758–68. doi: 10.1074/jbc.M113.534529. PubMed PMID: 24700463; PubMed Central PMCID: PMC4022850.

50. Worby CA, Mattoo S, Kruger RP, Corbeil LB, Koller A, Mendez JC, et al. The fic domain: regulation of cell signaling by adenylylation. Molecular cell. 2009; 34(1): 93–103. doi: 10.1016/j.molcel.2009.03.008. PubMed PMID: 19362538; PubMed Central PMCID: PMC2820730.

51. Paton AW, Srimanote P, Talbot UM, Wang H, Paton JC. A new family of potent AB(5) cytotoxins produced by Shiga toxigenic Escherichia coli. J Exp Med. 2004;200(1):35–46. doi: 10.1084/jem.20040392. PubMed PMID: 15226357; PubMed Central PMCID: PMCPMC2213318.

52. Paton AW, Beddoe T, Thorpe CM, Whisstock JC, Wilce MC, Rossjohn J, et al. AB5 subtilase cytotoxin inactivates the endoplasmic reticulum chaperone BiP. Nature. 2006;443(7111):548–52. doi: 10.1038/nature05124. PubMed PMID: 17024087.

53. Oakes SA, Papa FR. The role of endoplasmic reticulum stress in human pathology. Annu Rev Pathol. 2015; 10: 173–94. doi: 10.1146/annurev-pathol-012513-104649. PubMed PMID: 25387057.

54. Wang M, Kaufman RJ. The impact of the endoplasmic reticulum protein-folding environment on cancer development. Nat Rev Cancer. 2014; 14(9): 581–97. doi: 10.1038/nrc3800. PubMed PMID: 25145482.

55. Lee J, Ozcan U. Unfolded protein response signaling and metabolic diseases. J Biol Chem. 2014; 289(3): 1203–11. doi: 10.1074/jbc.R113.534743. PubMed PMID: 24324257; PubMed Central PMCID: PMC3894306.

56. Eletto D, Chevet E, Argon Y, Appenzeller-Herzog C. Redox controls UPR to control redox. J Cell Sci. 2014;127(Pt 17):3649–58. doi: 10.1242/jcs.153643. PubMed PMID: 25107370.

